# Profiling Human iPSC-Derived Sensory Neurons for Analgesic Drug Screening Using a Multi-Electrode Array

**DOI:** 10.1101/2024.11.18.623405

**Authors:** Christian Fofie Kuete, Rafael Granja-Vazquez, Vincent Truong, Patrick Walsh, Theodore Price, Swati Biswas, Gregory Dussor, Joseph Pancrazio, Benedict Kolber

**Author notes:** **Corresponding Author:** Benedict Kolber, 800 W. Campbell Road Richardson, TX 75080.

## Abstract

Chronic pain is a major global health issue, yet effective treatments are limited by poor translation from preclinical studies to humans. To address this, we developed a high-content screening (HCS) platform for analgesic discovery using hiPSC-derived nociceptors. These cells were cultured on multi-well micro-electrode arrays to monitor activity, achieving nearly 100% active electrodes by week two, maintaining stable activity for at least two weeks. After maturation (28 days), we exposed the nociceptors to various drugs, assessing their effects on neuronal activity, with excellent assay performance (Z’ values >0.5). Pharmacological tests showed responses to analgesic targets, including ion channels (Nav, Cav, Kv, TRPV1), neurotransmitter receptors (AMPAR, GABA-R), and kinase inhibitors (tyrosine, JAK1/2). Transcriptomic analysis confirmed the presence of these drug targets, although expression levels varied compared to primary human dorsal root ganglion cells. This HCS platform facilitates the rapid discovery of novel analgesics, reducing the risk of preclinical-to-human translation failure.

**Motivation:** Chronic pain affects approximately 1.5 billion people worldwide, yet effective treatments remain elusive. A significant barrier to progress in analgesic drug discovery is the limited translation of preclinical findings to human clinical outcomes. Traditional rodent models, although widely used, often fail to accurately predict human responses, while human primary tissues are limited by scarcity, technical difficulties, and ethical concerns. Recent advancements have identified human induced pluripotent stem cell (hiPSC)-derived nociceptors as promising alternatives; however, current differentiation protocols produce cells with inconsistent and physiologically questionable phenotypes.

To address these challenges, our study introduces a novel high-content screening (HCS) platform using hiPSC-derived nociceptors cultured on multi-well micro-electrode arrays (MEAs). The “Anatomic” protocol, used to generate these nociceptors, ensures cells with transcriptomic profiles closely matching human primary sensory neurons. Our platform achieves nearly 100% active electrode yield within two weeks and demonstrates sustained, stable activity over time. Additionally, robust Z’ factor analysis (exceeding 0.5) confirms the platform’s reliability, while pharmacological validation establishes the functional expression of critical analgesic targets. This innovative approach improves both the efficiency and clinical relevance of analgesic drug screening, potentially bridging the translational gap between preclinical studies and human clinical trials, and offering new hope for effective pain management.

## Introduction

Affecting nearly 1.5 billion people globally^1^, chronic pain profoundly impacts life quality and contributes to a higher incidence of mood disorders, particularly anxiety and depression^2^. This global crisis underscores the urgent need for new and effective pain treatments, especially given tolerance, dependence and addiction associated with opioid medications^3^. The traditional drug discovery process for analgesics is inefficient and prone to inaccuracies, in part because it relies on rodents and rodent cells, leading to untranslatable results^4^. For analgesic drug development targeting the peripheral nervous system, there is a need for more reliable screening platforms. Primary human tissues, though accurate, are often limited by scarcity, ethical concerns, and practical challenges, hindering their widespread use in screening^5^. To aid in the direct translation of drug discovery to humans, a robust and stable source of sensory neurons is needed to model the physiology and responsivity of the dorsal root ganglia (DRG).

The DRG are essential relay hubs for the transmission and modulation of nociceptive signals^6^. They contain the cell bodies of sensory neurons, including those specialized in detecting harmful or noxious stimuli. Those DRG nociceptors consist of small-to-medium diameter cells, which have either thinly myelinated (Aδ) or unmyelinated (C) axons^7^. Activated by various molecules like histamine, serotonin, and acetylcholine^8^, these neurons initiate nocifensive responses after peripheral injury. Nociceptors, particularly C fibers, are highly heterogeneous. Classically, these cells were divided into molecular sub-types based on expression of common markers. Peptidergic C nociceptors produce substance P, calcitonin gene-related peptide (CGRP) and pituitary adenylate-cyclase-activating polypeptide (PACAP), along with glutamate as neurotransmitters^9–12^. Non-peptidergic nociceptors were mainly known for expressing RET^13^, Mrgprd, Mrgpra3, and somatostatin^14^, for binding the isolectin B4 and for coexpressing the purinergic P2X_3_ receptor^15^. The release of nociceptor neurotransmitters in the dorsal horn of the spinal cord is modulated by several signals through Gi/o coupled receptors, such as GABA, opioid, cannabinoid, and adenosine receptors^16^. Nociceptors were also characterized by the presence of the tyrosine kinase receptors and the transient receptor potential (TRP) channels, including TRPV1 and TRPA1^17,18^. Additionally, they express a distinctive set of voltage-gated sodium, calcium, and potassium channels^19^. Recent studies have demonstrated additional heterogeneity through sequencing of both rodent and human DRG under both control and post-injury conditions^12,14,20–22^. The distinctions between species highlight the need to carefully consider the model of choice during analgesic drug screening with a clear need for more human-centric approaches.

While human immortalized cells provide some potential, their altered physiology from normal cells limits their utility for broad screening of analgesics. Human induced pluripotent stem cells (hiPSCs) present a promising alternative. These cells have the potential to revolutionize the identification and development of new therapeutic agents. At the core of this approach is the ability of hiPSCs to be derived directly from a patient’s own cells. This feature not only ensures a high degree of genetic and phenotypic similarity to the individual’s native cells but also captures the intricate complexity of human pathophysiology^5^, a level of detail that is often lost in traditional animal models or immortalized cell lines. The capacity to generate hiPSCs in large quantities further enhances their utility in drug discovery, addressing the long-standing issue of limited availability of primary human tissues and enabling the scale required for high-throughput screening of new drugs. Although there have been historical challenges in generating and maintaining hiPSC-derived nociceptors, recent work by Kalia et al.^23^ has successfully developed a differentiation process called the “Anatomic” protocol. This protocol produces nociceptor-like cultures with higher consistency than those obtained from the “Chambers” protocol^24^ and these cells are now commercially available as RealDRG™. These hiPSC nociceptors contain a diverse population of neuron-like cells which express many of the markers associated with nociceptors including Nav1.7, Nav1.8, and TRPV1 channels, reminiscent of primary human nociceptors^23^. Given these similarities to human cells, it is important to consider hiPSC nociceptors for their potential in high-content screening (HCS) for drugs.

HCS is an important component of current and future pain drug discovery and biomedical exploration, promoting the rapid testing of myriad chemical, genetic, or pharmacological manipulations by evaluating several features of neuron activity during drug application. Within the HCS, the multi-well Micro-Electrode Array (MEA) platform has emerged as a promising instrument. MEA platforms include arrays of electrodes within scalable culture plates. These systems allow for the non-invasive tracking of electrical impulses across heterogeneous cells^25,26^. The combination of MEA technology with hiPSC-derived nociceptors marks a significant leap forward, permitting the instantaneous observation of electrically excitable cells following drug application.

Although the MEA-hiPSC platform has emerged as a promising tool for drug discovery, practical implementation depends on the accuracy of the assay within the natural variability of hiPSC nociceptor excitability. Assessment of a platform’s effectiveness and reliability is achieved through the evaluation of the Z’ and/or robust Z’ factors^27,28^. The Z’ factor, a metric of assay performance when data come from a Gaussian distribution, offers insight into the assay’s capability to differentiate between active and inactive compounds with Z’ values generally considered excellent at 0.5 and ideal at 1^27^. The robust Z’ factor^28^ uses median-based analysis to account for data containing outliers. This is particularly valuable in the context of HCS coupled with hiPSC-derived nociceptors, as it accommodates the biological variability present in these cells.

Previously, we demonstrated the application of mouse DRG primary neurons to develop a MEA-based screening platform, achieving a robust Z’ factor of 0.61 with at least 25-50% of microelectrode contacts showing clear neural activity^28^. Here, we explore the hypothesis that using hiPSC nociceptors obtained from the “Anatomic” protocol could yield comparable or higher values. This approach was anticipated to broaden the screening scope and produce results with greater clinical relevance given the human origin of the Anatomic hIPSC nociceptors. The primary objective of our investigation was to ascertain the assay’s reliability by identifying the optimal seeding density that yields the most favorable Z’ or robust Z’ factors. Secondly, we sought to functionally characterize the hiPSC nociceptors in these MEAs through pharmacological probing, to identify pathways that could be targeted during novel analgesic screening. Overall, our results support the use of these cells for screening of nociceptor inhibitors agnostic to the precise molecular target, opening up the possibility of identifying novel chemical structures and targets in future *de novo* screening experiments.

## Results

### High active electrode yield achieved through optimized neuronal culture conditions

This experiment explored cell culture conditions, seeding techniques, and post-plating protocols to optimize active electrode yield (AEY) in hIPSC nociceptors derived using the Anatomic protocol (**Figure S1**). Electrode surfaces were pre-treated with Poly-L-Ornithine and coated with laminin-511 E8 to enhance neuronal attachment and growth. Sensory neurons were seeded onto 48-well and 96-well MEA plates at varying densities to study the impact on electrode coverage and AEY. For the 48-well MEA plate, densities ranged from 2K to 65K cells per well (**Figure 1**), while for the 96-well plate, densities were 15K, 35K, and 70K cells per well (**Figure S2**). Weekly monitoring of spontaneous activity was conducted, and cultures were maintained for up to 4 weeks to observe stable activity, including total spike counts (**Figure 1B, Figure S2A**), mean firing rate (**Figure 1C, Figure S2B**), and impedance (**Figure 1D, Figure S2C**).

**Figure 1:**
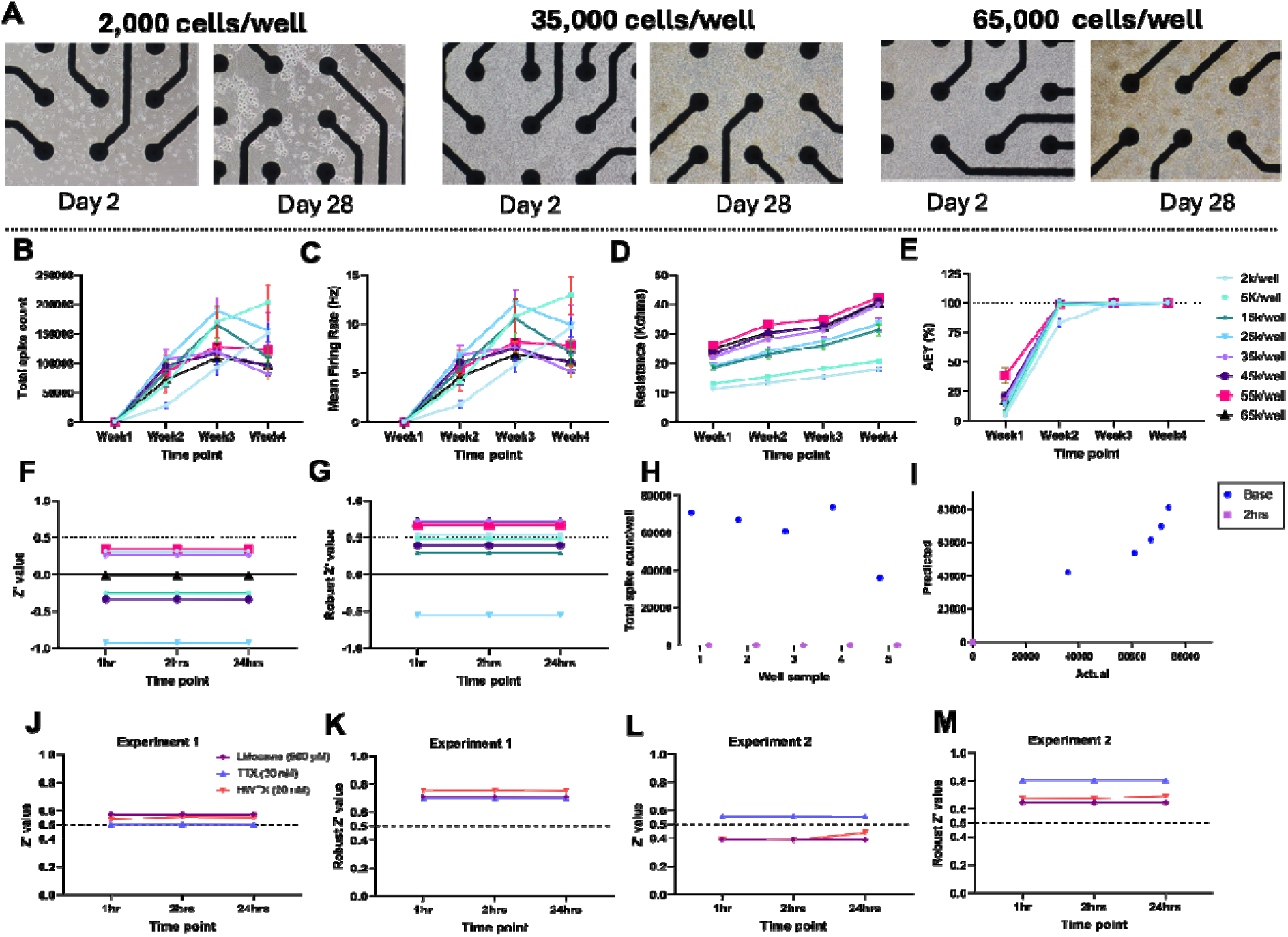
hiPSC nociceptors growth, neural activity, Z’ and Robust Z’ factors on a 48-well MEA plate. hiPSC nociceptors were cultured on a 48-well microelectrode array (MEA) plate for 4 weeks. **(A)** Representative micrographs of hiPSC nociceptors spot seeded at different cell densities (2,000, 35,000, and 65,000 cells/well) and two time points (day 2 and day 28). Across the four weeks of culture, cells were evaluated for **(B )** total spike count per well, **(C)** mean firing rate per well, **(D)** impedance, a measure of cell viability of hiPSC nociceptors, and **(E)** active electrode yield (AEY), indicating the percentage of electrodes successfully recording neural activity. At the 28 day time point, cells were challenged with inhibitors and the Z’ and robust Z’ factors were calculated to measure the quality of the assay. **(F)** Z’ and **(G)** robust Z’ factors were calculated respectively at different densities showing high quality with the robust Z’ factor at several seeding densities. **(H)** Scatter plots comparing well total spike count data before (blue) and 2 hours post-treatment with 500 µM lidocaine (pink) for a representative density of 35,000 cells/well. **(I)** Normal quantile-quantile (QQ) plot assessing the normality of data distribution before and after lidocaine treatment. Next, two independent experiments were conducted to evaluate the Z’ and robust Z’ of the assay to three neuronal inhibitors (lidocaine, tetrodotoxin (TTX), and huwentoxin (HWTX) for cells at 28 days in culture at 35,000 cells/well. **(J)** Z’ factor and **(K)** the robust Z’ values for different inhibitors in Experiment 1. **(L)** Z’ factor and **(M)** the robust Z’ values for different blockers in Experiment 2 using 35K/well. 0.5 dotted lines represents Z’ threshold for a “good” quality assay.

Results showed AEY reached 100% after 2 weeks across all densities, except for 2K cells per well (**Figure 1E**). In 96-well plates, AEY of ≥90% was achieved at 15K and 35K cells per well after 2 weeks (**Figure S2D**). At 70K cells per well, AEY reached ≥90% after 3 weeks, up from 73% at 2 weeks, indicating a longer culture time was needed for optimal AEY. This high AEY exceeded the 25-50% recommendation by Atmaramani et al^28^, highlighting the efficiency of the optimized conditions and techniques.

### MEA-hiPSC nociceptors platform reliability is validated by robust Z’ exceeding 0.5

The MEA-hiPSC nociceptor platform demonstrated reproducibility and sensitivity, with a robust Z’ factor exceeding 0.5, confirming its reliability in detecting changes in neuronal activity. Various cell densities in a 48-well format were tested with 500 µM lidocaine, which fully silenced cells without cytotoxicity, as shown by full recovery post-washout (**Figure S3A**). Initial per-electrode data revealed a non-normal distribution of spike rates, and log transformation was applied (**Figure S4**) but failed to fully normalize the data (**Table 1**). Only the 15K and 25K densities did not meet the classic Z’ factor threshold of 0.5 (**Figure S4E**). However, robust Z’ values post-log transformation exceeded 0.5 for all cell densities (**Figure S4F**).

**Table 1:**
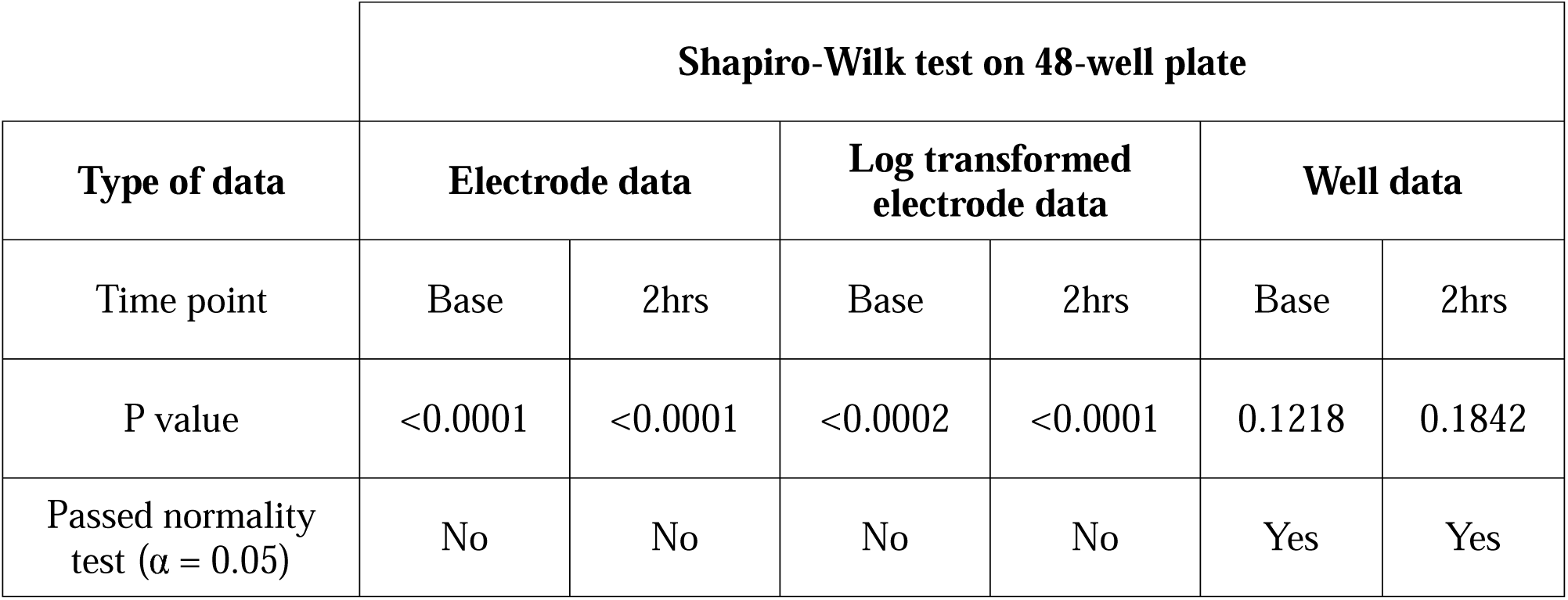
Distribution of data obtained from 48-well plate using Shapiro-Wilk test.

|  | Shapiro-Wilk test on 48-well plate |  |  |  |  |  |
| --- | --- | --- | --- | --- | --- | --- |
| Type of data | Electrode data |  | Log transformed electrode data |  | Well data |  |
| Time point | Base | 2hrs | Base | 2hrs | Base | 2hrs |
| P value | <0.0001 | <0.0001 | <0.0002 | <0.0001 | 0.1218 | 0.1842 |
| Passed normality test ($\alpha = 0.05$) | No | No | No | No | Yes | Yes |

At the well level, data followed a normal distribution (**Table 1**), eliminating the need for log transformation (**Figure 1**). Although none of the cell densities achieved classic Z’ values above 0.5 (**Figure 1F**), robust Z’ factors exceeded Z’ = 0.5 at 2K, 35K, 55K, and 65K cell densities (**Figure 1G**). Surprisingly, wells with 5K, 15K, and 25K cells did not reach the 0.5 threshold, unlike 2K wells. Based on these results, well-level data, which showed less variability than per-electrode data, was chosen for subsequent experiments.

A cell density of 35K cells/well was selected, and two confirmatory experiments using the robust Z’ factor were performed with three positive controls: lidocaine (500 µM), huwentoxin (HWTX, 20 nM), and tetrodotoxin (TTX, 30 nM). Both experiments yielded robust Z’ factors above 0.5 for all positive controls (**Figure 1K** and **1M**), while the classic Z’ factor did not consistently meet the threshold across replicates (**Figure 1J** and **1L**).

In 96-well MEA plates, cells treated with lidocaine (500 µM) achieved classic Z’ values of 0.5 at 15K and 70K cell densities in per-electrode analyses (**Figure S5A**, **S5C**) when data were Log transformed (**Figure S2E-S2H**) The Log robust Z’ factor exceeded 0.5 for all cell densities treated with lidocaine (**Figure S5D-F**). For TTX, only 15K and 70K cell densities reached a robust Z’ factor of 0.5 (**Figure S5D**), while huwentoxin-treated wells did not meet this threshold for any cell density. Per-well data in the 96-well plates followed an approximate normal distribution (p = 0.0473 at the 2-hour post-treatment point) (**Table S1**, **Figure S2I-J**). Z’ and robust Z’ factors calculated from per-well data revealed a robust Z’ value exceeding 0.5 only for the 15K cells/well condition (**Figure S5J**). Thus, the 15K cells/well density was selected for future experiments in 96-well MEA plates.

### TTX-sensitive and TTX-resistant voltage-Gated Sodium channels are functionally expressed in hiPSC nociceptors

After establishing the quality of the assay using the robust Z’ factor, we next sought to characterize the responses of hIPSC nociceptors in the context of known modulators of human primary nociceptors under baseline (37°C) and temperature ramp (37° to 42°C) conditions. Voltage-gated sodium channels (VGSCs) are crucial for initiating and propagating action potentials in DRG neurons. To assess their role in hiPSC nociceptors, we characterized TTX-sensitive (TTX-S) Nav1.7 and TTX-resistant (TTX-R) Nav1.8 channels using various inhibitors. TTX, at concentrations of 5-30 nM, completely silenced spontaneous activity at 15 nM in both 37°C and 42°C conditions (**Figure 2A-C**; **Table S2**), with peak inhibition within the first minute, sustained for 24 hours, and reversible within one hour after washout.

**Figure 2:**
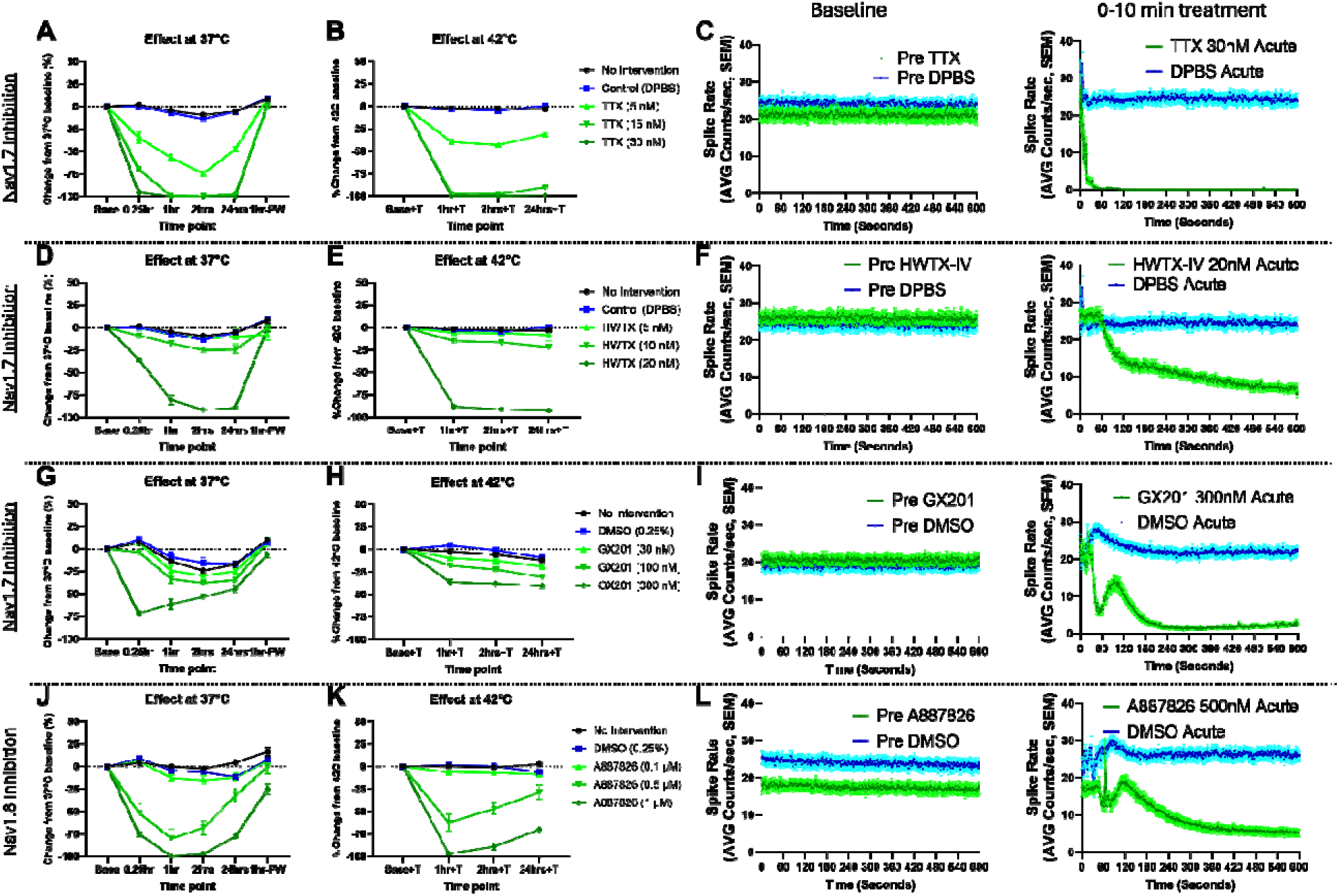
Inhibitory effects of sodium channel blockers on hiPSC nociceptor activity. The inhibitory effects of various sodium channel blockers on the activity of hiPSC nociceptors were assessed after 28 days of culture. Panels **A-C**, **D-F**, **G-I**, and **J-L** represent data obtained using tetrodotoxin (TTX, Nav1.7 blocker), huwentoxin IV (HWTX, Nav1.7 blocker), GX201 (Nav1.7 blocker), and A887826 (Nav1.8 blocker), respectively. For each blocker, the percentage inhibition of neuronal activity at 37°C and 47°C is shown (**A**, **B**, **D**, **E**, **G**, **H**, **J**, **K**) and was calculated relative to 37°C or 42°C baseline, respectively. Additionally, the spike rate of hiPSC nociceptors was monitored for 10 minutes before and after treatment with the respective blocker (**C**, **F**, **I**, **L**) at 37°C to evaluate acute effects. (See also Figure S6 and table S2)

Nav1.7 functionality was further confirmed using GX201 and Huwentoxin-IV. GX201 (300 nM) reduced activity by up to 71% at 37°C and 40% at 42°C, with recovery post-wash (**Figure 2D-F**). Huwentoxin-IV (20 nM) was more potent, reducing activity by 91% at 37°C and 92% at 42°C, with a gradual onset within the first minute and peak inhibition at one hour (**Figure 2G-I**). Full recovery followed washout.

For Nav1.8 (TTX-R), A887826 significantly reduced activity by 79% and 99% at 0.5 µM and 1 µM, respectively, at 37°C, and by 62% and 96% at 42°C (**Figure 2J-L**). Transcriptomic analysis revealed expression of SCN1A, SCN5A, SCN9A, and SCN10A, encoding Nav1.1, Nav1.5, Nav1.7, and Nav1.8 channels, respectively (**Figure S6**), confirming the presence of both TTX-S and TTX-R channels as targets for analgesic screening using hiPSC nociceptors.

### L- and T-type but not N-type calcium channels are potential targets for analgesic screening using hiPSC nociceptors

Voltage-gated calcium channels (VGCCs) are crucial for nociceptive signal transmission, with L-type, T-type, and N-type channels being the most relevant for peripheral analgesic screening. Mibefradil (T-type inhibitor) significantly reduced cell activity at 1 and 2 µM, decreasing by 40% and 89% at 37°C and by 28% and 77% at 42°C (**Figure 3A-C**; **Table S2**).

**Figure 3:**
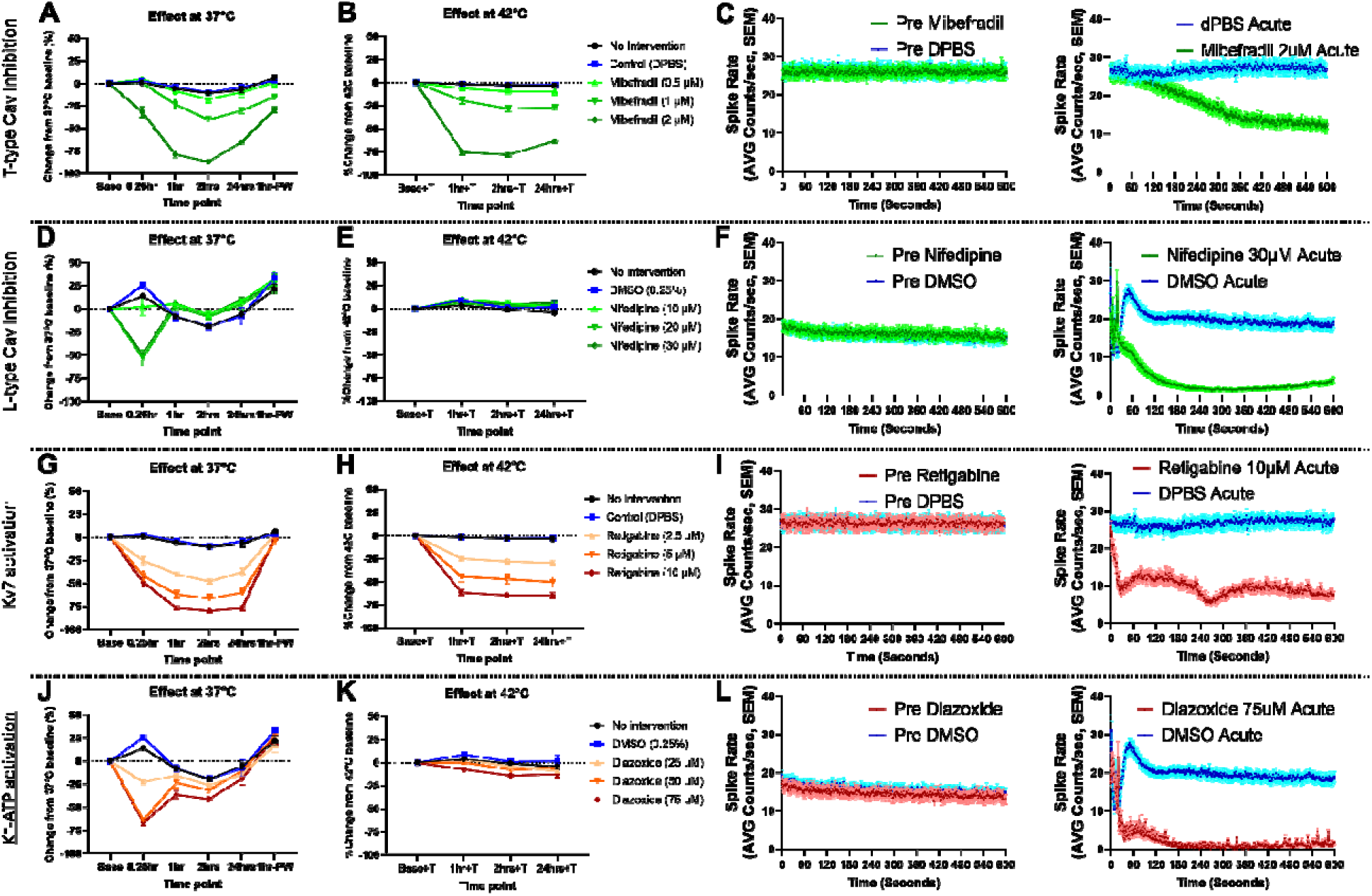
Effects of calcium channel blockers and potassium channel openers on hiPSC nociceptor activity. The inhibitory effects of these various drugs on the activity of hiPSC nociceptors were assessed after 28 days of culture. Panels **A-C** and **D-F** represent data obtained using mibefradil (T-type Ca^2+^ blocker) and nifedipine (L-type Ca^2+^ blocker) respectively. Panels **G-I** and **J-L** represent data obtained using retigabine (Kv7 opener) and diazoxide (K-ATP channel opener), respectively. For each blocker or activator, the percentage inhibition of neuronal activity at 37°C and 47°C is shown (**A**, **B**, **D**, **E**, **G**, **H**, **J**, **K**) and was calculated relative to 37°C or 42°C baseline, respectively. Additionally, the spike rate of hiPSC nociceptors was monitored for 10 minutes before and after treatment with the respective blocker (**C**, **F**, **I**, **L**) at 37°C to evaluate acute effects. (See also Figure S7, S8, S9 and table S2).

Nifedipine (L-type inhibitor) reduced activity by 49% and 52% at 20 and 30 µM, with a rapid onset and return to baseline within 1 hour at 37°C (**Figure 3D-F**). However, no effect wa observed at 42°C. ω-Conotoxin GVIA (N-type inhibitor) did not reduce activity at any concentration or temperature (**Figure S7A-C**). Transcriptomic analysis of hiPSC nociceptors showed strong expression of genes encoding L- and T-type calcium channels (**Figure S8**), identifying them as key targets for analgesic screening.

### Kv and Kir potassium channels are functionally expressed in hiPSC nociceptors

Potassium channels are essential for maintaining resting membrane potential and regulating neuronal excitability. This study explored voltage-gated potassium channels (Kv) and ATP-dependent potassium channels (Kir6). Retigabine (Kv agonist) at 2.5–10 µM significantly reduced hiPSC nociceptor activity by 47%–79% at 37°C and 29%–64% at 42°C, with effects starting within 1 minute (**Figure 3G-I**; **Table S2**). Diazoxide (Kir agonist) at 25–75 µM reduced activity by 22%–66%, peaking within 3 minutes but dissipating within an hour, with negligible effects at 42°C (**Figure 3J-L**). Transcriptomic analysis revealed strong expression of KCNQ2, KCNQ3, KCNQ4, and KCNJ11, encoding Kv7.2, Kv7.3, Kv7.4, and Kir6.2 channels (**Figure S9**), validating these channels as promising analgesic screening targets.

### AMPA/Kainate and TRPV1 receptors are functionally expressed in hiPSC nociceptors

Targeting glutamate-activated AMPA, kainate, and NMDA receptors offers a potential therapeutic strategy for pain management. DNQX, an AMPA/kainate receptor antagonist, significantly reduced hiPSC nociceptor activity by 48%–84% at 37°C, with effects diminishing by 60% after 24 hours, while efficacy was weaker at 42°C (18%–32% reduction) (**Figure 4A-C**; **Table S2**). Transcriptomic analysis confirmed the expression of AMPA (GRIA1-4) and kainate (GRIK1-5) receptor subunits (**Figure S10**). However, NMDA antagonist APV showed no effect (data not shown).

**Figure 4:**
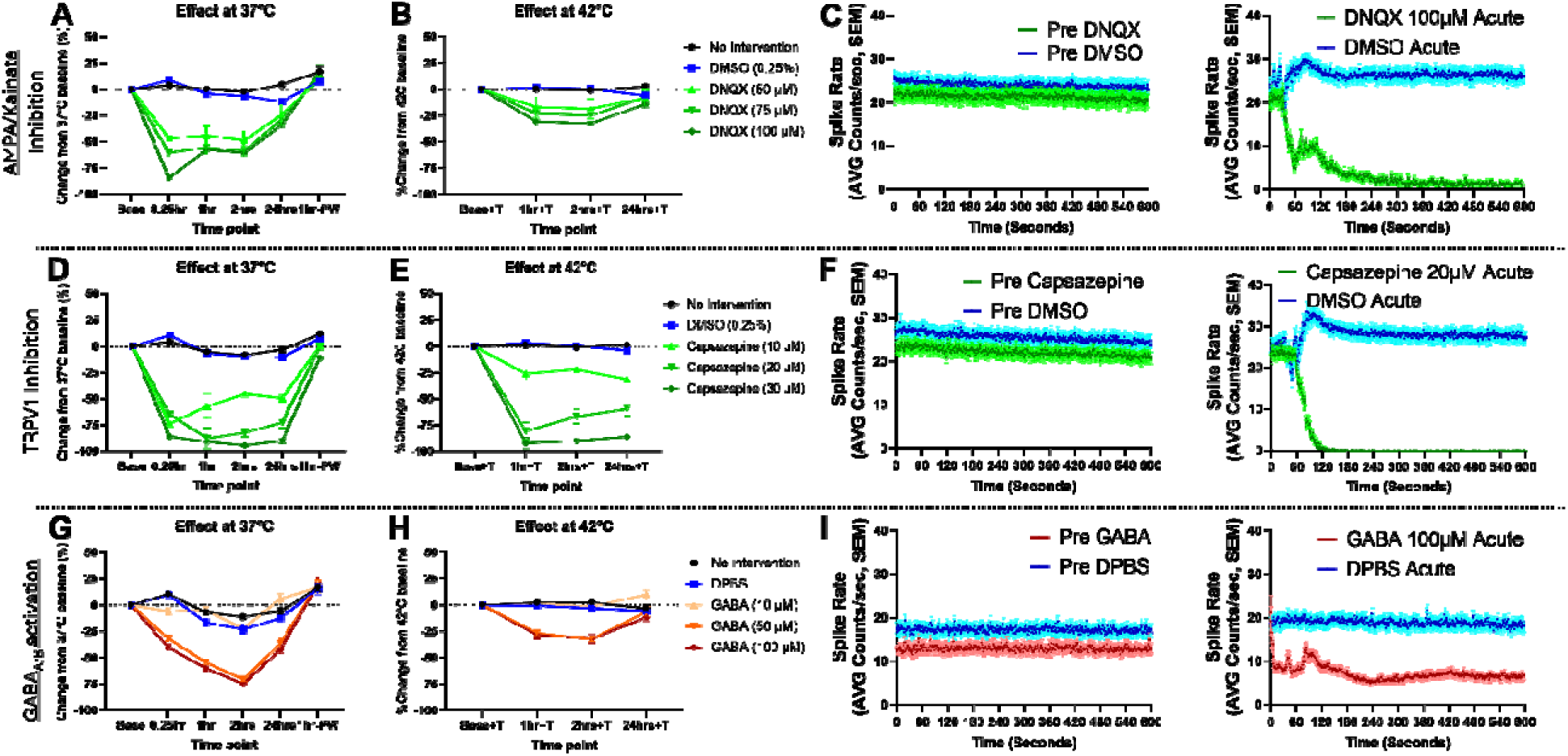
Effects of AMPA/kainite receptor antagonist, TRPV1 channel blocker and GABA receptor activation on hiPSC nociceptor activity. The inhibition of glutamate AMPA/Kainite receptors and TRPV1 channels, and the activation of GABA receptors, respectively on the activity of hiPSC nociceptors were assessed after 28 days of culture. Panels **A-C**, **D-F** and **G-I**, represent data obtained using DNQX (AMPA/Kainate antagonist), capsazepine (TRPV1 antagonist), and GABA (*pan*-GABA receptor agonist) respectively. For each drug, the percentage inhibition of neuronal activity at 37°C and 47°C is shown (**A, B, D, E, G, H**) and was calculated relative to 37°C or 42°C baseline, respectively. Additionally, the spike rate of hiPSC nociceptors was monitored for 10 minutes before and after treatment with the respective blocker (**C, F, I**) at 37°C to evaluate acute effects. (See also Figure S10, S11, S12 and table S2).

For TRPV1 receptors, capsazepine significantly reduced nociceptor activity at 37°C by 73%–94% and at 42°C by 31%–91%, with rapid onset and return to baseline within 1 hour post washout (**Figure 4D-F**; **Table S2**). High TRPV1 expression in hiPSC nociceptors was detected, though lower than in primary human DRG (**Figure S11**), highlighting AMPA/kainate and TRPV1 receptors as viable targets for analgesic screening using this platform.

### GABA but not mu-opioid receptors are functionally expressed in hiPSC nociceptors

GABA and mu-opioid receptors (MOR) play key roles in modulating pain at the DRG-CNS synapse. In this study, GABA significantly reduced hiPSC nociceptor activity at 50 and 100 µM concentrations by 70% and 74% at 37°C, with effects peaking at 2 hours and a milder reduction (31%) observed at 42°C (**Figure 4G-I**; **Table S2**). Transcriptomic analysis confirmed the presence of GABAA and GABAB receptors (**Figure S12**).

In contrast, DAMGO (MOR agonist) at 1-10 µM had no significant effect on cell activity at either temperature (**Figure S7D-F**), despite robust OPRM1 gene expression encoding MOR (**Figure S13**), suggesting limited functional relevance of MOR when screening for analgesic using this MEA-hiPSC nociceptors platform.

### Kinases as targets when screening for analgesic using hiPSC nociceptors

Various kinases involved in nociception were evaluated using inhibitors to determin their potential as target. Dasatinib, a broad-spectrum kinase inhibitor, significantly reduced hiPSC nociceptor activity at 5-20 µM concentrations, with reductions of 64%–91% at 37°C and 22%–84% at 42°C, showing a rapid onset within 2 minutes (**Figure 5A-C**; **Table S2**).

**Figure 5:**
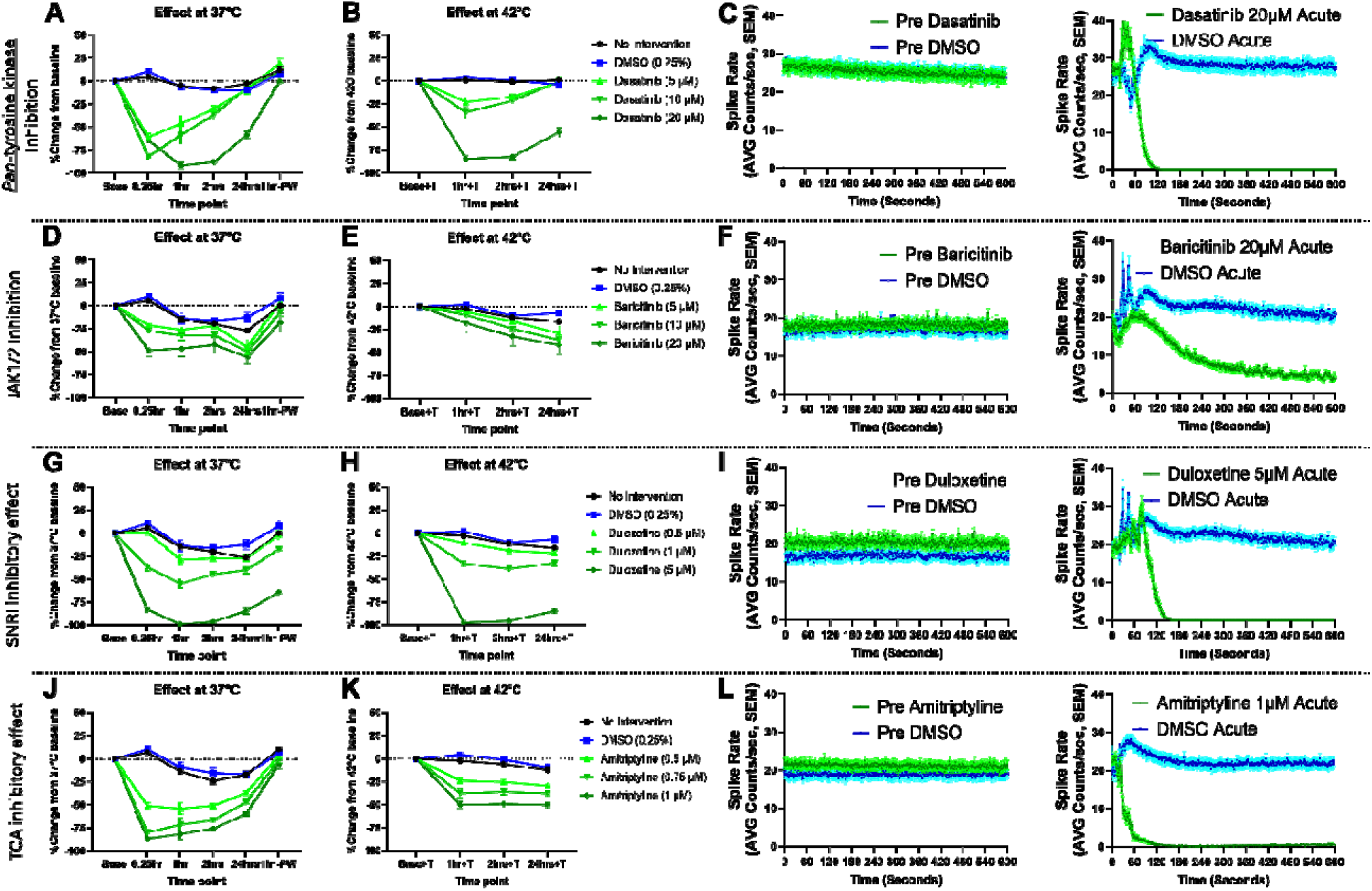
Inhibitory effects of kinase inhibitors, TCA and SNRI on hiPSC nociceptors. The inhibitory impact of various kinase inhibitors on the activity of hiPSC nociceptors following a 28-day culture period. Panels **A-C,** and **D-F** represent data obtained using dasatinib (multiple tyrosine kinase inhibitor) and baricitinib (JAK1/2 inhibitor), respectively. Panels **G-I** and **J-L** represent data obtained using duloxetine (serotonin and norepinephrine reuptake inhibitor) and amitriptyline (tricyclic antidepressant), respectively. For each inhibitor, the percentage inhibition of neuronal activity at 37°C and 47°C is shown (**A, B, D, E, G, H, J, K**) and was calculated relative to 37°C or 42°C baseline, respectively. Additionally, the spike rate of hiPSC nociceptors was monitored for 10 minutes before and after treatment with the respective blocker (**C, F, I, L**) at 37°C to evaluate acute effects. (See also Figure S14 and table S2).

Baricitinib, targeting JAK1/2, decreased activity by 44%–55% at 37°C and by 29%–41% at 42°C, peaking at 24 hours (**Figure 5D-F**). Conversely, MNK1/2 inhibitor eFT-508 showed no significant effect up to 1 µM, regardless of temperature (**Figure S7G-H**; **Table S2**). Transcriptomic data confirmed high expression of these kinases in hiPSC nociceptors (**Figure S14**), validating their potential as therapeutic targets for analgesic screening.

### Validating the screening assay using clinical off-labeled drugs for neuropathic pain

Duloxetine, a selective norepinephrine reuptake inhibitor, and amitriptyline, a tricyclic antidepressant, were evaluated for their ability to suppress hiPSC nociceptor activity. Duloxetine demonstrated concentration-dependent inhibition at 37°C, reducing cell activity by 28%, 64%, and 98% at 0.5, 1, and 5 µM, respectively, with a rapid onset within 3 minutes (**Figure 5G-I**; **Table S2**). This effect was sustained for an hour, and at 42°C, significant reductions of 38% and 97% were observed, with the highest concentration maintaining suppression even after washout.

Amitriptyline also effectively reduced activity at 37°C by 53%, 79%, and 86% at 0.5, 0.75, and 1 µM, respectively, with peak inhibition occurring within the first minute (**Figure 5J-L**; **Table S2**). However, its efficacy was diminished at 42°C, showing reduced inhibition rates of 29%, 37%, and 50% at the same concentrations, highlighting temperature-dependent effects on its analgesic potential.

## Discussion

Pain research has long faced limitations due to the reliance on traditional *in vitro* and *in vivo* rodent models, prompting the need for faster and more efficient methods in analgesic drug discovery. The combination of human iPSC nociceptor cultures with MEA technology offers a promising solution to complete early screening in human cells. iPSC-derived nociceptors provide a physiologically relevant *in vitro* system that closely replicates the complex neural responses to injury, addressing both ethical concerns and the variability inherent in animal models. Meanwhile, MEA technology enables high-content screening by monitoring multiple neurons simultaneously, delivering real-time, quantitative data on drug efficacy.

Despite these advantages, MEA technology presents its own challenges that can affect data reliability. Although MEAs effectively capture electrical signals from neurons, issues such as electrode degradation and inconsistent signal quality can lead to unreliable results. A critical metric for assessing MEA performance is the active electrode yield (AEY), which is the percentage of microelectrode sites exhibiting at least one spike per minute. Atmaramani et al^28^ determined that an AEY of 25-50% is necessary to produce reliable data from mouse DRG neurons in Axion MEA setups. In our study, we achieved nearly 100% AEY within two weeks, which may be attributed to factors such as surface chemistry, seeding density, and advances in the Axion plate reader technology. The Axion PEDOT electrodes were coated with poly-L-ornithine (PLO), enhancing cell attachment, as shown by Ge et al^29,30^ who found that PLO promotes better proliferation, migration, and differentiation of neural stem cells compared to Poly-L-lysine and fibronectin. Laminin 511E8 was also used to coat the wells, providing more effective long-term cell growth and differentiation support than Matrigel^31,32^.

To further explore factors affecting AEY, the study examined different seeding densities, ranging from 2,000 to 70,000 cells per well, using both classic and spot-seeding techniques. After 28 days, the AEY remained consistently high (∼100%) across all densities and techniques, suggesting that seeding density had little effect on the ability of hiPSC nociceptors to form functional electrical connections. This is likely due to the pseudo-unipolar structure of DRG neurons and their ability to generate bidirectional action potentials^33^, which enhances neuron-electrode interactions and contributes to the high AEY.

However, a high AEY alone does not guarantee reliable drug screening outcomes. Given the complexity of nociceptor signaling pathways, platforms must consistently identify active compounds, and robust quality control measures are crucial for ensuring data integrity. The Z’ factor, introduced by Zhang et al^27^ and refined by Atmaramani et al^28^, is an essential metric for evaluating assay quality. Z’ factors between 0.5 and 1.0 indicate excellent assays, while values below 0 suggest overlap between control signals, making the assays unreliable. In our study, log transformation was applied to electrode spike count data to account for non-normal distributions, resulting in excellent Z’ factors. This non-normality likely stemmed from the heterogeneity of the cell population^34^.

The study focused on total spike counts across wells as the primary metric for analyzing positive control agents. Unlike electrode-level data, the well-based data followed a normal distribution without the need for log transformation. West^35^ cautioned that applying log transformation when zero observations exceed 2% of the dataset can introduce bias. Our analysis revealed that only certain plating densities achieved robust Z’ factors (≥ 0.5) in 48- and 96-well plates, with better results observed in the spot-seeding technique used for the 48-well plate. Although a minimum effective density of 2,000 cells per well was identified, we opted for 35,000 cells per well in subsequent experiments to avoid overestimating drug potency and ensure accurate screening results.

Ultimately, validating the use of the Axion MEA with Anatomic’s hiPSC nociceptors required confirming that these cells responded to known analgesics in a manner similar to human DRG cells. The overall pharmacologic screening results, displayed in **Figure 6**, support the reliability of this system for analgesic drug discovery. Nav channels are crucial for the rapid depolarization phase of the action potential in excitable cells^36^. The nine Nav channel subtypes display distinct biophysical properties and tissue-specific expression patterns^37^, reflecting their diverse pharmacology and physiological roles. Notably, mutations in the SCN9A gene, encoding the Nav1.7 channel, provide compelling evidence for its essential contribution to pain perception suggesting that Nav1.7 may be a promising target for the development of novel analgesics^38^. To assess the role of Nav1.7 in our hiPSC nociceptors, we used three structurally and mechanistically distinct Nav1.7 blockers. The first blocker, tetrodotoxin (TTX), potently silenced neuronal activity at 30 μM. TTX acts by binding to the external selectivity filter of tetrodotoxin-sensitive voltage-gated sodium channels (TTX-S VGSCs), thereby inhibiting sodium influx and action potential generation^39^. While TTX is commonly used as a Nav1.7 blocker at 30 µM on nociceptors, it exhibits broader selectivity. Studies have shown that most VGSCs are sensitive to low TTX concentrations (<30 nM), while Nav1.5, 1.8, and 1.9 require higher concentrations (>1 μM) for blockade^37^. Given that adult sensory neurons express Nav1.1, 1.6, 1.7, 1.8, and 1.9^40^, the complete silencing observed with TTX likely reflects combined contributions from the inhibition of Nav1.1 and Nav1.6 alongside Nav1.7. This hypothesis is supported by the incomplete silencing observed with huwentoxin-IV (HWTX) at 20 nM, a concentration known to be specific for Nav1.7 at this concentration^41^. Unlike TTX, which does not directly affect channel gating mechanisms, HWTX inhibits the activation of sodium channels by trapping the voltage sensor of domain II of the site 4 in the inward, closed configuration^41^. The third specific Nav1.7 used was GX201, which inhibits the voltage-dependent gating of domain IV of the channel^42^. This mechanism of action differs from that of huwentoxin-IV, which likely explains the observed disparity in the inhibitory effect. While GX201 exhibited inhibitory nociceptor neural activity, its potency even at 300 nM was lower compared to HWTX. Additionally, elevated temperature markedly reduced the impact of GX201. Among TTX-resistant channels, Nav1.8, a voltage-gated sodium channel (VSGC) encoded by the SCN10A gene, is of particular interest due to its specific expression in nociceptive neurons, highlighting its critical role in pain signaling pathways^43^. To investigate its function, we used the selective Nav1.8 blocker A-88782623 at concentrations up to 1 µM^23^. This blocker produced a near-complete inhibition (almost 100%) of channel activity at both physiological (37°C) and elevated (42°C) temperatures. This assay provides robust evidence for its efficacy in identifying analgesic drugs that target and inhibit Nav1.7 and Nav1.8 sodium channels.

**Figure 6:**
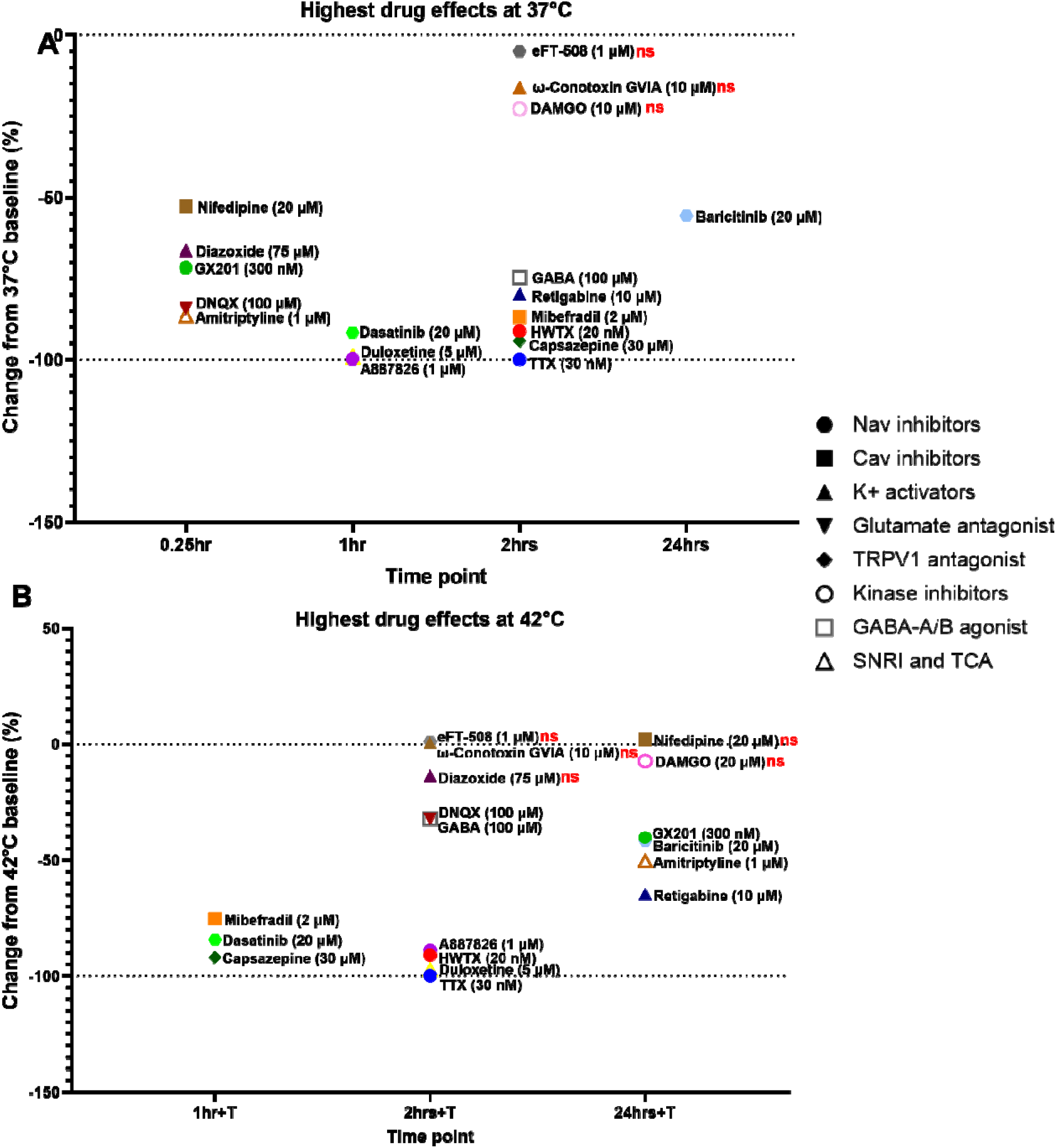
Functional channels, receptors, and kinases in hiPSC nociceptors at 37°C and during 42° C heat ramps. Each time point indicates peak drug efficacy at **(A)** 37°C and **(B)** during 42° C heat ramps.

The next channel family that was characterized were the calcium channels (Cav). Cav channels are crucial in various physiological processes, including neurotransmitter release, muscle contraction, and gene expression^44^. Their role in nociceptor signaling has gained significant attention in recent years. Cav channels are composed of a pore-forming α1 subunit and auxiliary subunits (β, α2δ, and γ). The α1 subunit, which determines the channel’s biophysical and pharmacological properties, is divided into several types: Cav1 (L-type), Cav2 (P/Q-type, N-type, R-type), and Cav3 (T-type)^45^. We investigated the functionality of each type using specific blockers. Nifedipine, a selective L-type Cav channel blocker, significantly reduced sensory neuron activity with a maximum effect at 20 μM. This effect was transient, lasting up to 15 minutes. The observed potency of nifedipine aligns with the findings of Newberry et al.^46^, who reported a 67% reduction in activity of embryonic rat DRG neurons at 10 μM. Interestingly, in our study, 10 μM nifedipine did not elicit any effect. This discrepancy might be attributed to species-specific differences in Cav channel expression or sensitivity. T-type voltage-gated calcium channels also play a role in pain signaling. Matsunami et al.^47^ demonstrated that mibefradil, a non-selective T-type Ca² channel blocker, effectively prevented and reduced visceral nociceptive behavior. To investigate the contribution of T-type Cav channels in our system, we tested mibefradil. Notably, 2 μM of this drug significantly reduced DRG activity by up to 80%. Among Cav2 channels, N-type channels are probably the most relevant to pain signaling within DRG neurons, since P/Q-type channels primarily contribute to descending facilitatory systems, and R-type channels are predominantly localized to the cell soma and proximal neuronal dendrites^48^. We used ω-conotoxin GVIA to investigate the functionality of N-type Cav channels in DRG. Despite using ω-conotoxin GVIA known to be more efficacious than ziconotide, the FDA-approved N-type channel inhibitor for pain^49^, at a high concentration of 10 μM, we did not observe any inhibitory activity on DRG sensory neurons. This lack of effect could be attributed to the predominantly synaptic localization of N-type Cav channels within DRG neurons^49,50^. Moreover, N-type calcium channels are predominantly found in the peripheral nervous system where they play a pivotal role in neurotransmitter release at autonomic and sensory nerve terminals^51,52^. Since our cultured hiPSCs likely lack functional synapses, ω-conotoxin GVIA was therefore irrelevant in this assay^47^. Taken together, this assay demonstrates its utility in screening for modulators of L-type and T-type calcium channels while being less suitable for investigating N-type channel activity.

DRG neurons express a diverse repertoire of voltage-gated K channels, including Kv1, Kv3, Kv4, Kv7, KCa, and Kir subfamilies^53,54^. These channels are critical regulators of neuronal excitability by controlling K efflux, influencing membrane repolarization and hyperpolarization. Dysfunction of K channels is implicated in the development and persistence of neuropathic pain^55^. In this study, we investigated the functional effects of retigabine and diazoxide respectively on KCNQ/Kv7 and ATP-sensitive K /Kir6.2 channels in hiPSC nociceptors. As expected, application of 10 µM retigabine significantly reduced neuronal activity by nearly 80% and the effect lasted at least 24 hours. Retigabine’s mechanism of action involves potentiating Kv7.2-7.5 channels, which are known to maintain resting membrane potential and regulate neuronal excitability^56^. Diazoxide exhibited a transient inhibitory effect, reaching a maximum at 50 µM. This inhibition decayed over time, with the effect persisting for up to 2 hours at 75 µM. These findings align with Kawano et al.^57^, who demonstrated that diazoxide (at 100 µM), but not the Kir6.1 inhibitor glybenclamide, hyperpolarized the resting membrane potential of axotomized neurons in hyperalgesic rats following spared nerve injury (SNL). Further supporting the role of Kir6.2 channels in neuropathic pain, Luu et al.^58^ showed that mice lacking SUR1 subunits of K_ATP_ channels displayed mechanical hypersensitivity, which was alleviated by diazoxide (300 µg/paw) only within the first 20 minutes of treatment. In this study, we focused on investigating only two potassium channels, but other K channels could also represent additional potential targets for drug discovery alongside those examined here.

Peripheral glutamate receptors have emerged as a promising target for novel analgesic therapies due to their role in modulating nociceptive transmission. DRG neurons express mRNA or immunoreactivity for both ionotropic and metabotropic glutamate receptors^59^. Ultrastructure studies have provided evidence for the transport of AMPA and NMDA receptors into peripheral processes of these neurons^60^. Additionally, DRG neurons innervating the colon respond to NMDA with increased intracellular calcium levels^61^. In this study, we investigated the functionality of AMPA and kainate receptors using DNQX. Treatment with this drug resulted in a significant and sustained decrease in DRG cell activity for up to 2 hours. The 50 µM concentration of DNQX used in our study exceeded the reported IC ^62^, though similar concentrations have been used in other research^63,64^. Conversely, the NMDA antagonist APV did not elicit any observable effect (*data not shown*). These findings suggest a crucial role for AMPA and kainate receptors in the hiPSC activity, highlighting their potential for screening with these cells. Further studies are warranted to explore NMDA receptor functionality, including extended maturation as genotyping has revealed mRNA expression of these receptors in these hiPSC nociceptors.

Transient receptor potential (TRP) channels are cation channels expressed in various tissues by both excitable and non-excitable cells. These channels play a crucial role in modulating the driving force for the influx of cations like Na^+^, K^+^, Ca^2+^, and Mg^2+65^. Several TRP subfamilies, including TRPV, TRPM, TRPA, and TRPC, are known to be involved in nociception and pain perception^66^. Notably, TRPV1 displays a characteristic of metabolic coupling to various neural receptors that recognize molecules inducing pain and itch sensations^67^. This functional coupling suggests that targeting TRPV1 activity presents a valuable and rational therapeutic strategy. In this study, we evaluated TRPV1 function using capsazepine at concentrations up to 30 µM. Capsazepine significantly reduced hiPSC nociceptor activity at both 37°C and during 42°C temperature ramps for up to 24 hours post-treatment. This finding may appear to contradict established literature, as TRPV1 activation typically occurs at temperatures ranging from 40°C to 45°C^68^. However, capsazepine possesses documented off-target effects, including modulation of Janus kinase (JAK) signaling and downregulation of LPS-induced NF-κB activation^69^. This is of particular importance as baricitinib, a specific JAK1/2 inhibitor, significantly reduced hiPSC nociceptor activity at 37°C but exhibited minimal effect at 42°C. This observation suggests that capsazepine’s action at 37°C may stem from its pleiotropic effects, while its activity at 42°C is more attributable to TRPV1 channel inhibition. An alternative explanation is the hiPSC nociceptors may exist in a “pre-sensitized” state as one might find after an injury. In this context, some of the TRPV1 receptors may be active at the 37°C baseline state even prior to the temperature ramp. This pre-sensitization hypothesis will be explored in future studies. Overall, these findings demonstrate the assay’s potential for effectively screening TRPV1 antagonists.

Kinase inhibition has emerged as a promising therapeutic strategy in the field of analgesia, particularly for chronic pain management. Kinases are a class of enzymes responsible for phosphorylating protein substrates^70^, a critical step in various cellular signaling pathways, including those involved in pain perception^71^. The human kinome classifies conventional protein kinases into eight groups based on sequence homology and functional similarities. These groups include tyrosine kinases, CMGC, AGC, CAMK, CK1, and atypical/other kinases (e.g., IKK, mTOR, MNK)^72^. In this study, we investigated the effects of dasatinib (a multi-tyrosine kinase inhibitor), baricitinib (a JAK1/2 inhibitor targeting cytokine signaling), and eFT-508 (an MNK inhibitor, a recently identified target in pain management)^73^. Both dasatinib and baricitinib significantly reduced hiPSC nociceptor activity, with dasatinib’s effect persisting even at 42°C. Interestingly, MNK inhibition did not elicit a decrease in sensory neuron activity. This may be attributed to the potential immaturity of MNK function in the cells at the time of investigation since mRNA for MNK1/2 are expressed in these cells. Nonetheless, the findings with dasatinib and baricitinib align with those of Appel et al.^74^ and Makabe et al.^75^, who demonstrated reduced pain in cancer-induced bone pain and arthritis-induced pain models, respectively. Therefore, the assay platform we have developed demonstrates usefulness in screening some kinase inhibitors for their potential to modulate pain pathways. However, it is important to acknowledge that the age of the cells may constitute a limitation to the spectrum of kinase inhibitors that can be investigated.

Mu-opioid receptor (MOR) agonists represent a crucial class of medications valued for their potent analgesic effects. Mechanisms for this property could involve DRG as Moy et al^76^ reported that about 50% of human DRG neurons express functional “MOR-like” receptors. To investigate MOR activity in our assay, we used DAMGO, a specific MOR agonist, at concentrations up to 10 µM^77^. DAMGO treatment failed to demonstrate any inhibitory effect on the activity of hiPSC nociceptors. This finding aligns with aforementioned experiment regarding the inhibition of N-type calcium channels. It has been established that G protein-coupled opioid receptors suppress neuronal activity by inhibiting voltage-gated N-type calcium channels^78^. Given the likely absence of synapses in our culture, DAMGO’s potential to reduce cell activity through synaptic modulation was precluded. These observations align with the work of Newberry et al.^46^, who demonstrated that morphine at concentrations up to 5 µM failed to induce a decrease in activity in rat embryonic stem cell-derived DRG neurons.

Gamma-aminobutyric acid (GABA) receptors are critical modulators of pain signaling within the DRG^79^. These receptors are categorized into two main types: GABA-A and GABA-B, both of which are expressed in DRG neurons^79^. In the central nervous system (CNS), GABA typically acts as an inhibitory neurotransmitter^80^ by causing hyperpolarization, making neurons less likely to fire action potentials. This occurs because GABA opens chloride channels, allowing Cl^−^ ions to flow into the neuron, increasing its negative charge. In contrast, in dorsal root ganglia (DRG) neurons, GABA causes depolarization^81^ due to a higher intracellular Cl^−^ concentration, leading to a net outflow of Cl^−^ ions. Despite this depolarization, GABA in DRG neurons often reduces excitability through a shunting effect, preventing the generation of action potentials^82^. Studies by Du et al.^82^ demonstrated that *in vivo* administration of GABA or a GABA reuptake inhibitor to sensory ganglia significantly reduced acute peripheral pain and alleviated neuropathic and inflammatory pain in rats. However, Valeyev et al.^83^ reported that GABA receptors in cultured human DRG neurons exhibit pharmacological properties distinct from those observed in rodents. We therefore challenged the cells with GABA at concentrations up to 100 µM and observed a significant inhibitory effect. This finding corroborate previous studies and suggests that the assay can be used to screen for GABAergic agonists. However, further investigation is necessary to determine the specific GABA receptor subtype(s) targeted by the assay.

Finally, to confirm the overall capacity of this platform to screen for relevant clinical analgesics, we challenged the cells with a TCA and a SNRI. Although these drugs were not initially developed for pain management, they are increasingly recognized as first-line therapies for neuropathic pain^84^. In this study, we investigated the effects of amitriptyline (TCA) and duloxetine (SNRI) on DRG activity. Both drugs significantly reduced hIPSC nociceptor activity. Amitriptyline’s analgesic effects are known to involve modulation of voltage-gated sodium and potassium channels^85^, and calcium channels^86^. Similarly, duloxetine has been shown to block specific Nav^87,88^ and Kv^89^ channels. Given our assay’s sensitivity to Nav, Kv, and Cav channel inhibitors (**Figure 6**), the observed reductions in DRG activity by amitriptyline and duloxetine are likely attributable to their modulation of these ion channels. These findings therefore support the use of this assay for screening ion channel inhibitors with potential relevance to pain pathways.

Several studies, including ours, have developed platforms using rodent models or hiPSC-derived neurons for drug screening. Recently, another model was introduced using hiPSC-derived neurons and astrocytes on MEAs to screen potential analgesics^90^. However, concerns about the phenotypic fidelity of the model arise due to the formation of a networked, synaptically-connected culture which is not typical of DRG cultures, and therefore could affect the accuracy of drug screening results. The co-culture component of that model offers a significant exploration into the hiPSC nociceptor landscape. Additionally, a comparison could be drawn with another model that used a modified “Chambers” method for hiPSC differentiation. In a limited MEA experiment performed after 2 weeks in culture, data demonstrated that DAMGO at 5 µM could inhibit hiPSCs following ATP activation—an effect that was not observed at concentrations up to 10 µM in our experiments. However, the absence of an evaluation of other potential nociceptor modulators in that study makes a thorough comparison challenging.^91^.

### Limitations and Conclusion

In summary, we report in this study the development of a novel HCS platform for analgesic drug discovery that uses hiPSC-derived nociceptors differentiated using the Anatomic protocol. There are some limitations of this assay system as highlighted by our data. The assay relies upon microelectrodes that detect external spikes as a surrogate for action potential firing in neurons. Any mechanism of action for analgesia that could be detected, for example, in patch clamp electrophysiology, but that does not result in a decrease in action potential firing in cells would be missed in this assay. Furthermore, while the cells used in this study share transcriptional similarity to human primary nociceptors^23^ compared to rodent primary nociceptors, not every target with a transcriptional signature may be functional in the cells. In the present study, this appears to be true for both the targets MNK and Ca_v_2.2, as the nociceptors failed to respond with a decrease in action potential firing to eFT-508 and ω-conotoxin GVIA, respectively.

Furthermore, these cells are active at baseline and could even be considered to be in a pre-sensitized state as one would find after injury. They are not however at a maximum with respect to firing rate since activity can be further enhanced through heat ramp application. For drug screening purposes, this seeming biological drawback may be an advantage because the cells can be tested “as is” without manipulation. Nonetheless, this assay overcomes limitations of conventional HCS methods by incorporating functionally relevant and scalable human cells. We successfully validated the platform’s functionality by demonstrating its response to several, but not all, established analgesic drugs, inflammatory stimuli, and by confirming expression of key cellular targets associated with pain signaling. Taken together, this innovative approach has the potential to significantly accelerate the identification of safe and clinically effective pain medications through both targeted screening for molecules with a proposed mechanism as well as unbiased target-agnostic screening of small molecule and natural product libraries.

## STAR Methods

### General

All reagents and resources utilized can be found in **Table S3**. When possible, experiments were completed blinded to experimental treatment and are indicated as such in below methods sections. In all cases, technical replicates and pilot experiments were utilized to determine concentration ranges for all drugs and compounds.

### hiPSC-derived nociceptor generation

Commercially available RealDRG (hiPSC nociceptors) were manufactured as previously described^23^. In brief, immature post-mitotic sensory neurons were generated using a scaled-up version of Anatomic’s Senso-DM kit (Anatomic, CAT#1010). These cells were cryopreserved at a concentration of 1 million viable neurons per vial before being shipped on dry ice to The University of Texas at Dallas. All cells used in this study were derived from the human iPSC line ANAT001, which were reprogrammed from the cord blood of a healthy adult female using episomal plasmids. Nociceptors derived from this hiPSC line have been previously characterized in a number of different assays and applications^23,92–95^. The cells used were from lot number SN308214, which passed standard quality control measures including neuronal purity > 95%, verified cell number per vial, post-thaw viability > 70%, and sterility.

### hiPSC nociceptor transcriptomics

The time course expression of individual genes in hiPSC-nociceptors were referenced from Anatomic’s RealDRGene App v2.0 (https://anatomicincorporated.shinyapps.io/realdrgene/). This database was generated using Anatomic hiPSC line ANAT001 and four weekly time points from three lots of hiPSC nociceptors manufactured from this line as previously described by Truong et al^96^. In short, hiPSC-nociceptors sequencing libraries were generated using an Illumina TruSeq Stranded mRNA Library Prep Kit. Equimolar quantities from each library were sequenced on a NextSeq 550 High Output Kit v2.5 (75 cycles) at approximately 30 million single-end reads per sample. The primary human DRG data set referenced in the application was previously published^97^.

### Seeding and maintenance of hiPSC nociceptors

To functionally evaluate hiPSC nociceptors, multi-well Axion 48-well or 96-well MEAs were used. For 48-well MEA plates, the following procedures were used. Prior to initiating the culture, the MEA was prepared by coating with Poly-L-Ornithine (0.01%) and incubating overnight at room temperature. After three washes with sterile deionized water, each well received iMatrix-511 SILK coating (1:50 dilution with dPBS without calcium and magnesium) and was incubated for three hours at 37°C. A single tube of hiPSC nociceptors was thawed by gradually increasing the temperature from -80°C to 25°C over a period of approximately 1 minute at 37°C. Following thawing, the cells were washed using DMEM/F12, centrifuged at 300g for 4 minutes and subsequently resuspended in 1 mL of Anatomic Senso-MM complete growth medium. After counting the cells, they were centrifuged once more, and the pellet was resuspended to obtain the desired cell concentration in a volume of Senso-MM medium. Following the removal of excess iMatrix-511 from each well, cells were spot-seeded in the center of each electrode array at a density range of 2,500 to 75,000 cells per well (**Figure 1A**). This was done in a total volume of 5 µL and the cells were then incubated for 20 minutes to facilitate cell attachment. Then, in each well, a total volume of 400 µL of Anatomic Senso-MM was added slowly. Media exchanges (50% of 400 µL) were conducted the next day and every other day, with cells maintained at 37°C in 5% CO . Overall, media was changed three times per week (Monday, Wednesday, Friday) with 50% media exchanges. For 96-well MEA plates, all plate preparation conditions were identical to that described above for 48-well MEA plates except for the final seeding step. Instead of spot seeding, cells were seeded across the entire well at 15,000, 35,000 and 70,000 cells/well with a volume of 75 µL.

### Active Electrode Yield (AEY)

In the core AEY experiment, cells were spot-seeded at varying densities, ranging from 2,000 to 65,000 cells per 5 µL in 48-well MEA plates or well-seeded at densities of 15,000, 35,000, and 70,000 cells per 75 µL in 96-well MEA plates. They were maintained for four weeks as described above. MEA electrophysiological data were recorded using an Axion Maestro Pro and AxIS software at a 12.5 kHz sampling rate, applying a single-pole Butterworth bandpass filter (300-5000 Hz). Individual spikes were detected from continuous voltage recordings exceeding an adaptive ± 5.5σRMS threshold. Recordings were done once a week for four weeks at 37°C for 15 minutes. Data were always recorded 48 hours after a media change to avoid media-associated changes in excitability. For quantitative analysis, electrodes were classified as active if they produced a minimum average of one spike per minute across an entire 15- minute recording session. The plotted AEY represent the percentage of active electrodes out of 16 available per well in the 48-well format or out of 8 in the 96-well MEA. This experiment was done unblinded to cell density plating.

### Z’ and robust Z’ factors

During the 4^th^ week of cell culture, the baseline activity of hiPSC nociceptors in the MEA plates containing different cell densities was recorded. Lidocaine (500 µM) or a Nav channel blocker were added each to five out of six wells per density, while the 6th well was treated with vehicle (dPBS). Recordings were taken immediately after treatment for 15 minutes and followed by additional recordings at 1 hour, 2 hours, and 24 hours post-treatment at 37°C. Subsequently, the media in each well was completely replaced, and another recording was taken at least 1 hour later. The results obtained at the 1, 2, and 24-hour time points were used to calculate Z’ and robust Z’ factors using both electrode-based and well-based metrics. Electrode metrics focused on the individual performance of each electrode, measuring the number of spikes recorded by a single electrode in a 15-minute window. In contrast, well metrics looked at the overall activity of the well, capturing the total number of spikes recorded by all 16 electrodes within the same 15-minute period. When there was evidence that the data are not normally distributed, a log_10_ transformation was applied. For confirmation purposes, the experiment was repeated twice using a cell density of 35,000 cells/well. In these replicate experiments, cells were challenged with two additional Nav channel blockers, TTX (30 nM) and Huwentoxin IV (20 nM), with 12 wells per treatment and dPBS as the vehicle control. More detailed data analysis and statistics data can be found below in the Statistics section. This experiment was done unblinded to treatment condition.

### Pharmacological probing to phenotype hiPSC nociceptors

Initially, compounds were prepared by dissolving them in dimethyl sulfoxide (DMSO) or dPBS without calcium or magnesium ions. These solutions were then diluted with dPBS to achieve a stock concentration 80 times higher than the desired final concentration. Subsequently, the compounds were diluted 1:80 directly into the plate well, using 5 µL of the compound for every 400 µL of medium. During the experiments, DMSO at 0.25% was used as the final concentration. Under unsensitized conditions, pharmacological agents were tested across nine “molecular families,” including voltage-gated ion channel modulators, G-protein-coupled receptor (GPCR) agonists, kinase inhibitors, and atypical analgesics. The agents tested belonged to the following categories and concentration ranges were derived from literature followed by pilot testing (*data not shown*) in these cells:

| Compound Name | Target/Mechanism | Concentration Range |
| --- | --- | --- |
| Huwentoxin IV | Nav1.7 inhibitor | 5-20 nM<sup>28,41,98,99</sup> |
| TTX | Nav inhibitor | 5-30 nM<sup>99,100</sup> |
| GX201 | Nav1.7-selective blocker | 30-300 nM |
| A-887826 | Nav1.8-selective blocker | 0.1-1 μM<sup>23,101</sup> |
| Retigabine | KCNQ1-5/Kv7 opener | 2.5-10 μM<sup>56,102,103</sup> |
| Diazoxide | K<sup>+</sup>-ATP activator | 25-75 μM<sup>57,104</sup> |
| Nifedipine | L-type calcium channel blocker | 10-30 μM<sup>105-108</sup> |
| ω-Conotoxin GVIA | N-type calcium channel blocker | 0.1-10 μM<sup>108-110</sup> |
| Mibefradil | T-type calcium channel blocker | 0.5-2 μM<sup>107,111</sup> |
| GABA | GABA-A/B agonist | 10-100 μM<sup>83,112</sup> |
| eFT-508 | MNK inhibitor | 25-1000 nM<sup>113,114</sup> |
| Dasatinib | Pan-tyrosine kinase inhibitor | 5-20 μM<sup>115</sup> |
| Baricitinib | Janus kinase, JAK1/2 inhibitor | 5-20 μM<sup>116</sup> |
| DNQX | AMPA/kainate antagonist | 50-100 μM<sup>63,64</sup> |
| D-(-)APV | NMDA antagonist | 25-100 μM |
| Capsazepine | TRPV1 antagonist | 10-30 μM<sup>117-119</sup> |
| Duloxetine | Atypical SNRI | 0.5-5 μM<sup>46</sup> |
| Amitriptyline | Atypical TCA | 0.5-1 μM<sup>46,120</sup> |
| DAMGO | Mu-opioid agonist | 1-10 μM<sup>76,121</sup> |

Recordings were always started 24 hours after a media change to avoid media-associated excitability changes. Baseline activity was measured for 15 minutes at both 37°C and 42°C temperature ramps. Following this, recordings were taken immediately after drug treatment for 15 minutes. Subsequent recordings were made at 1 hour, 2 hours, and 24 hours post-treatment. Except for the 0.25-hour time point, all recordings were conducted at both 37°C and 42°C temperature ramps. After the 24-hour recording, the wells were washed, and a final measurement was taken at least one hour later at 37°C. At each time point, the total spike count per well was normalized to the corresponding baseline (37°C or 42°C temperature ramp). In this experiment, drug concentrations were pseudo-randomly assigned to experimental groups (n=6/group) in a blinded manner.

### Statistical methods for data analysis

Spontaneous and evoked activity data were extracted from Axion spike (.spk) files using Axion Biosystems’ Neural Metric Tool, with a minimum firing rate criterion of 1 spike per minute. Data are presented as mean ± SEM, calculated using the total spike count per well.

#### Z’ Analysis

Z’ was calculated using two methods.

First, in order to calculate the “classic” Z’ factor, the following formula from Zhang et al.^27^ was used:

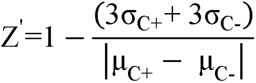

μ_c+_ and μ_c-_ represent the average values (means) of the positive and negative controls, respectively. σ_c+_ and σ_c-_ represent the standard deviations of the positive and negative controls, respectively.

Second, when using the robust Z’ factor, the following formula from Atmaramani et al^28^ was used:

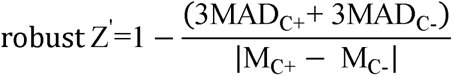

*MAD_c_*_+_ and *MAD_c_*_−_ refers to the median average deviation of the positive and negative controls and *M_c_*_+_ and *M_c_*_−_ refer to the median of the positive and negative controls, respectively. Classic and robust Z’ calculations assume a normal distribution. Therefore, for non-normal data, we applied a log_10_ transformation. At baseline, all electrodes showed >1 electrode spiking. However, after treatment, several electrodes demonstrated full inhibition (spikes = 0). To avoid log transformation with zero values during log transformation, we added 1 to each electrode’s total spike for all time points before the transformation. The normality test was performed using the Shapiro-Wilk test with the significance level (alpha) set at 0.05.

#### General analysis and statistics

Percent changes from baseline were calculated relative to the corresponding temperature baseline (either 37°C static or 37-42°C temperature ramp). All percent (%) changes from baseline were calculated using the following formula:

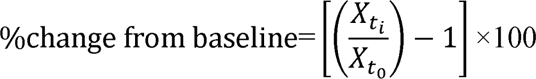

Where:

*X*: represents a data point.

*t*_0_: represents the baseline time point (before drug administration).

*t*_i_: represents one of the following time points: 0.25 hours, 1 hour, 2 hours, 24 hours, or 1 hour post-washout (1hr-PW).

In the pharmacological probing experiments, a two-way ANOVA with repeated measures was used to compare each positive control group with its corresponding vehicle control (either dPBS or DMSO). Dunnett’s post-hoc test was used for multiple comparisons within each positive control group. All graphs were created in Graphpad Prism (v10.0).

#### Rate Histograms

Files collected during the simultaneous addition of the compounds underwent further processing to extract temporal features of the firing rate. First, the .spk file was converted in the Data Export Tool (Axion Biosystems) and processed in NeuroExplorer from the .nex file format. Rate histograms were generated at spike counts per 1 second bins over the duration of the recording. Each compound is shown during two timepoints, the first 10 minutes prior to its addition during baseline and then for 10 minutes while it was being added to the well. Both sessions were collected at 37°C. For color consistency across our graphs, traces from agonists are shown in red, inhibitors are shown in green, and vehicles are shown in blue.

## Acknowledgements

This work was supported by NIH NCCIH IGNITE Grants R61-AT011938 and R33-AT011938 (BJK, GD, JP).

## Author contributions

**Conceptualization**: BLK, GD, JP; **Methodology:** BJK, RGV, JP, CFK, SB; **Investigation:** CFK, RGV, BJK; **Writing – Original Draft:** CFK, BJK, RGV; **Writing – Review & Editing:** TP, SB, GD, JP, BJK; **Funding Acquisition:** GD, JP, BJK; **Resources:** TP, VT, PW; **Supervision:** GD, JP, BJK.

## Declaration of interests

PW and VT are shareholders and employees of Anatomic Incorporated. hiPSC nociceptors “RealDRG” and associated reagents were provided by Anatomic for project through co-investigator status on R61/R33 award.

## Supplemental information

Document S1. Figures S1 – S14

Document S2. Table S1 – S3

## Figure Legends

**Figure S1: Overall methodology for technique.**

**Figure S2: Electrode and well data distribution of hiPSC nociceptors on 96-well MEA.** hiPSC nociceptors were cultured at varying densities on a 96-well MEA plate for four weeks. **(A)** Total spike count per well over 28 days; **(B)** Mean firing rate per well over 28 days; **(C) I**mpedance, a measure of cell viability of hiPSC nociceptors; **(D)** active electrode yield (AEY), indicating the percentage of electrodes successfully recording neural activity. Electrode and well total spike count served as the primary metric. **(E)** Scatter plot comparing electrode spike count data before (blue) and 2 hours post-treatment with 30 nM TTX (pink) for a representative density of 15,000 cells/well. **(F)** Normal quantile-quantile (QQ) plot assessing the normality of data distribution before and after TTX treatment. **(G)** Scatter plot of Log(x+1) transformed data for the same representative density as in panel A. **(H)** Normal QQ plot evaluating the normality of the Log(x+1) transformed data. **(I)** Scatter plot comparing well total spike count data before (blue) and 2 hours post-treatment with 30 nM TTX (pink) for a same density as in panel A. **(J)** Normal quantile-quantile (QQ) plot assessing the normality of data distribution before and after TTX treatment.

**Figure S3: Concentration-response inhibitory curves for lidocaine (A), tetrodotoxin (B), and huwentoxin-IV (C) on hiPSC-nociceptor activity.** These data were used to determine Z’ and robust Z’ factors. “PW” indicates post-wash measurements.

**Figure S4: Electrode data distribution, Z’, and robust Z’ factors of hiPSC nociceptors on 48-well MEA.** hiPSC nociceptors were cultured at varying densities on a 48-MEA plate for four weeks. Electrode total spike count served as the primary metric. **(A)** Scatter plot comparing electrode spike count data before (blue) and 2 hours post-treatment with 500 µM lidocaine (pink) for a representative density of 35,000 cells/well. **(B)** Normal quantile-quantile (QQ) plot assessing the normality of data distribution before and after lidocaine treatment. **(C)** Scatter plot of Log(x+1) transformed data for the same representative density as in panel A. **(D)** Normal QQ plot evaluating the normality of the Log(x+1) transformed data. **(E)** Z’ factor values calculated for Log(x+1) transformed data across different hiPSC nociceptor densities. 0.5 dotted line represents threshold for a “good” quality assay. **(F)** Robust Z’ factor values calculated for Log(x+1) transformed data across different hiPSC-DRG cell densities.

**Figure S5: Z’-factor analysis of hiPSC nociceptors cultured on 96-well MEA plates.** hiPSC-derived nociceptors were cultured at three densities (15,000, 35,000, and 70,000 cells/well) on 96-well microelectrode array (MEA) plates for four weeks. Total spike counts were determined for each electrode and each well. Classic and robust Z’-factor values were calculated based on spike count data from both electrode-level (log-transformed) and well-level analyses. (**A, D**) Classic and robust Z’-factors, respectively, for electrode-level analysis at a cell density of 15,000 cells/well. (**B, E**) Classic and robust Z’-factors, respectively, for electrode-level analysis at a cell density of 35,000 cells/well. (**C, F**) Classic and robust Z’-factors, respectively, for electrode-level analysis at a cell density of 70,000 cells/well. (**G, J**) Classic and robust Z’-factors, respectively, for well-level analysis at a cell density of 15,000 cells/well. (**H, K**) Classic and robust Z’-factors, respectively, for well-level analysis at a cell density of 35,000 cells/well. (**I, L**) Classic and robust Z’-factors, respectively, for well-level analysis at a cell density of 70,000 cells/well.

**Figure S6: Gene expression profile of sodium channels in hiPSCs, hiPSC nociceptors and hDRG**. (**A**) expression of SCN1A which encodes for Nav1.1. (**B**) expression of SCN5A encodes for Nav1.5. (**C**) expression of SCN9A which encodes for Nav1.7. (**D**) expression of SCN10A which encodes for Nav1.8. hiPSC are human iPSCs line ANAT001 (Anatomic hiPSC Female Line#1), which were reprogrammed from the cord blood of a healthy female using episomal plasmids. hiPSC-nociceptors are nociceptors manufactured from hiPSC Female Line#1. hDRG are primary human DRG that was previously published^96,97^. Gene expression is typically considered to be present when Rlog values exceed 6.

**Figure S7: Effects of N-type Ca^2+^ inhibitor, mu-opioid receptor activation and MNK1/2 inhibitor on hiPSC nociceptor activity.** The inhibitory effects of these drugs were assessed after 28 days of culture. Panels **A-C**, **D-F**, and **G-I** represent data obtained using ω-conotoxin GVIA (N-type Ca^2+^ inhibitor), DAMGO (mu-opioid agonist), and eFT-508 (MNK1/2 inhibitor) respectively. For each drug, the percentage inhibition of neuronal activity at 37°C and 47°C is shown (**A, B, D, E, G, H**) and was calculated relative to 37°C or 42°C baseline, respectively. Additionally, the spike rate of hiPSC nociceptors was monitored for 10 minutes before and after treatment with the respective blocker (**C, F, I**) at 37°C. (See also Figure S7, S12, S13 and table S2).

**Figure S8: Gene expression profile of calcium channels in hiPSC, hiPSC nociceptors and hDRG**. (**A)** CACNA1C which encodes for Cav1.2 (L-type). (**B**) CACNA1D which encodes for Cav1.3 (L-type). (**C**) CACNA1A which encodes for Cav2.1 (N-type). (**D**) CACNA1G which encodes for Cav3.1 (T-type). (**E**) CACNA1H which encodes for Cav3.2 (T-type). (**F**) CACNA1I which encodes for Cav3.3 (T-type). hiPSC are human iPSCs line ANAT001(Anatomic hiPSC Female Line#1), which were reprogrammed from the cord blood of a healthy female using episomal plasmids. hiPSC-nociceptors are nociceptors manufactured from hiPSC Female Line#1. hDRG are primary human DRG that was previously published^96,97^. Gene expression is typically considered to be present when Rlog values exceed 6.

**Figure S9: Gene expression profile of potassium channels in hiPSC, hiPSC-DRG and hDRG**. (**A**) KCNQ2 which encodes for Kv7.2. (**B**) KCNQ3 which encodes for Kv7.3. (**C**) KCNQ4 which encodes for Kv7.4. (**D**) KCNJ11 which encodes for Kir6.2/ATP-K^+^ . hiPSC are human iPSCs line ANAT001(Anatomic hiPSC Female Line#1), which were reprogrammed from the cord blood of a healthy female using episomal plasmids. hiPSC-nociceptors are nociceptors manufactured from hiPSC Female Line#1. hDRG are primary human DRG that was previously published^96,97^. Gene expression is typically considered to be present when Rlog values exceed 6.

**Figure S10: Gene expression profile of AMPA and kainate receptors in hiPSC, hiPSC nociceptors and hDRG**. (**A-D**) GRIA1, GRIA2, GRIA3, GRIA4, encodes for AMPA receptor subunits. (**E-I**) GRIK1, GRIK2, GRIK3, GRIK4 and GRIK5 encodes for kainate receptor subunits. hiPSC are human iPSCs line ANAT001(Anatomic hiPSC Female Line#1), which were reprogrammed from the cord blood of a healthy female using episomal plasmids. hiPSC-nociceptors are nociceptors manufactured from hiPSC Female Line#1. hDRG are primary human DRG that was previously published^96,97^. Gene expression is typically considered to be present when Rlog values exceed 6.

**Figure S11: Gene expression profile of TRPV1 channel in hiPSC, hiPSC nociceptors and hDRG**. TRPV1 gene encode TRPV1 channel. hiPSC are human iPSCs line ANAT001(Anatomic hiPSC Female Line#1), which were reprogrammed from the cord blood of a healthy female using episomal plasmids. hiPSC-nociceptors are nociceptors manufactured from hiPSC Female Line#1. hDRG are primary human DRG that was previously published^96,97^. Gene expression is typically considered to be present when Rlog values exceed 6.

**Figure S12: Gene expression profile of GABA receptors in hiPSC, hiPSC nociceptors and hDRG**. (**A-H**) GABRA2, GABRA3, GABRA5, GABRB3, GABRG2, GABRG3, GABRE and GABRQ which encode for GABA-A receptor subunits. (**I**) GABBR1 and (**J**) GABBR2 encode for GABA-B receptor subunits. hiPSC are human iPSCs line ANAT001(Anatomic hiPSC Female Line#1), which were reprogrammed from the cord blood of a healthy female using episomal plasmids. hiPSC-nociceptors are nociceptors manufactured from hiPSC Female Line#1. hDRG are primary human DRG that was previously published^96,97^. Gene expression is typically considered to be present when Rlog values exceed 6.

**Figure S13: Gene expression profile of mu-opioid receptor in hiPSC, hiPSC nociceptors and hDRG**. OPRM1 encodes for mu-opioid receptor. hiPSC are human iPSCs line ANAT001(Anatomic hiPSC Female Line#1), which were reprogrammed from the cord blood of a healthy female using episomal plasmids. hiPSC-nociceptors are nociceptors manufactured from hiPSC Female Line#1. hDRG are primary human DRG that was previously published^96,97^. Gene expression is typically considered to be present when Rlog values exceed 6.

**Figure S14: Gene expression profile of various kinases in hiPSC, hiPSC nociceptors and hDRG**. (**A-B**) MKNK1 and MKNK2 which encode for MAP kinase-interacting serine/threonine-protein kinase 1 and 2 (MNK1/2) respectively. (**D-E**) JAK1 and JAK2 which encode for Janus kinase 1 and 2 respectively. (**C, F, G**) Scr, ABL and EFNB2 encode for sarcoma tyrosine kinase, Abelson kinase and Ephrin B2 receptor tyrosine kinase, respectively. hiPSC are human iPSCs line ANAT001(Anatomic hiPSC Female Line#1), which were reprogrammed from the cord blood of a healthy female using episomal plasmids. hiPSC-nociceptors are nociceptors manufactured from hiPSC Female Line#1. hDRG are primary human DRG that was previously published^96,97^. Gene expression is typically considered to be present when Rlog values exceed 6.

## Notes

### Competing Interest Statement

Patrick Walsh and Vincent Truong are shareholders and employees of Anatomic Incorporated. hiPSC nociceptors -RealDRG- and associated reagents were provided by Anatomic for project through co-investigator status on R61/R33 award.

