## Supplementary figures and images for "Profiling Human iPSC-Derived Sensory Neurons for Analgesic Drug Screening Using a Multi-Electrode Array"

### Supplemental Files Figure S1-S14

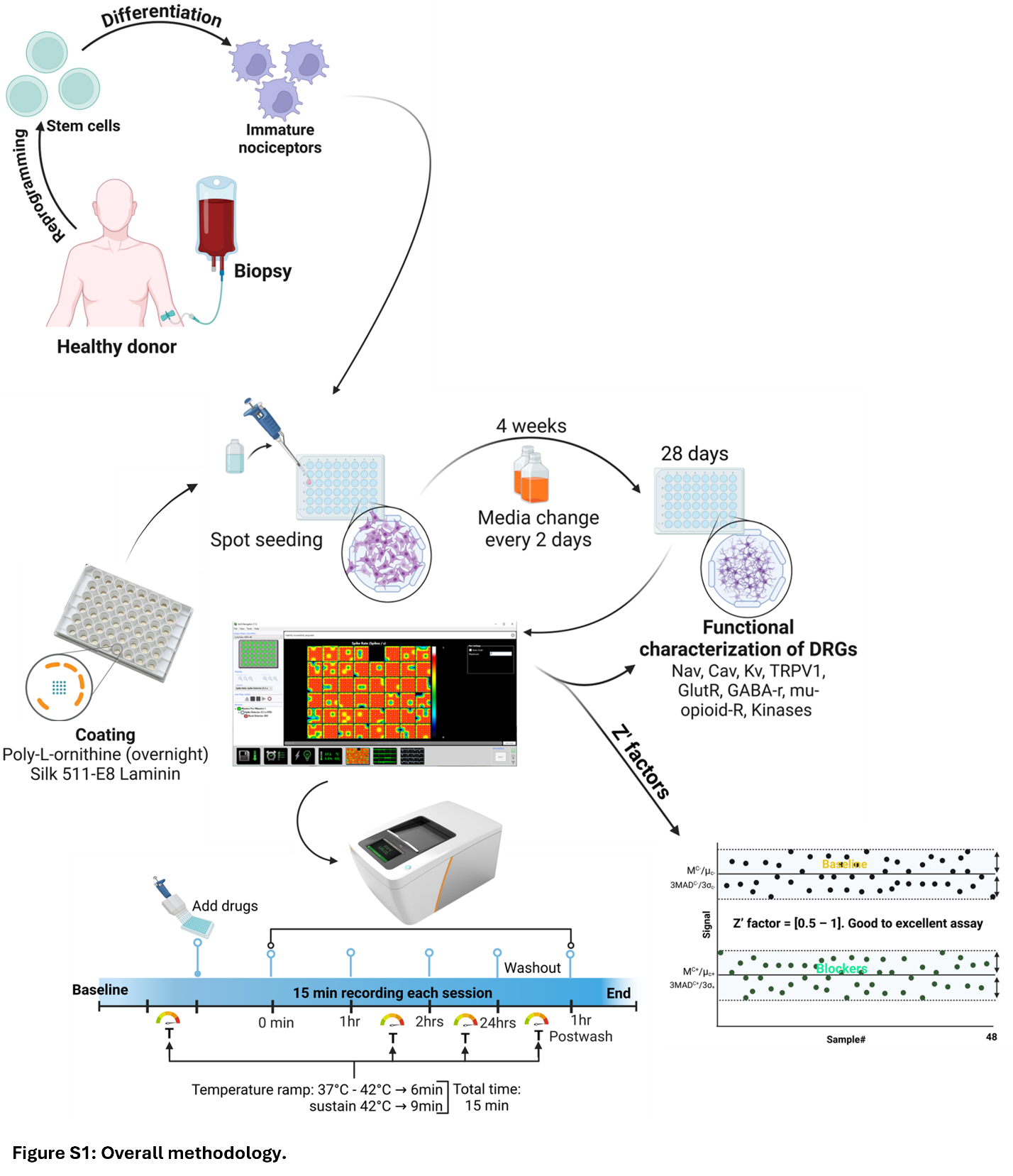

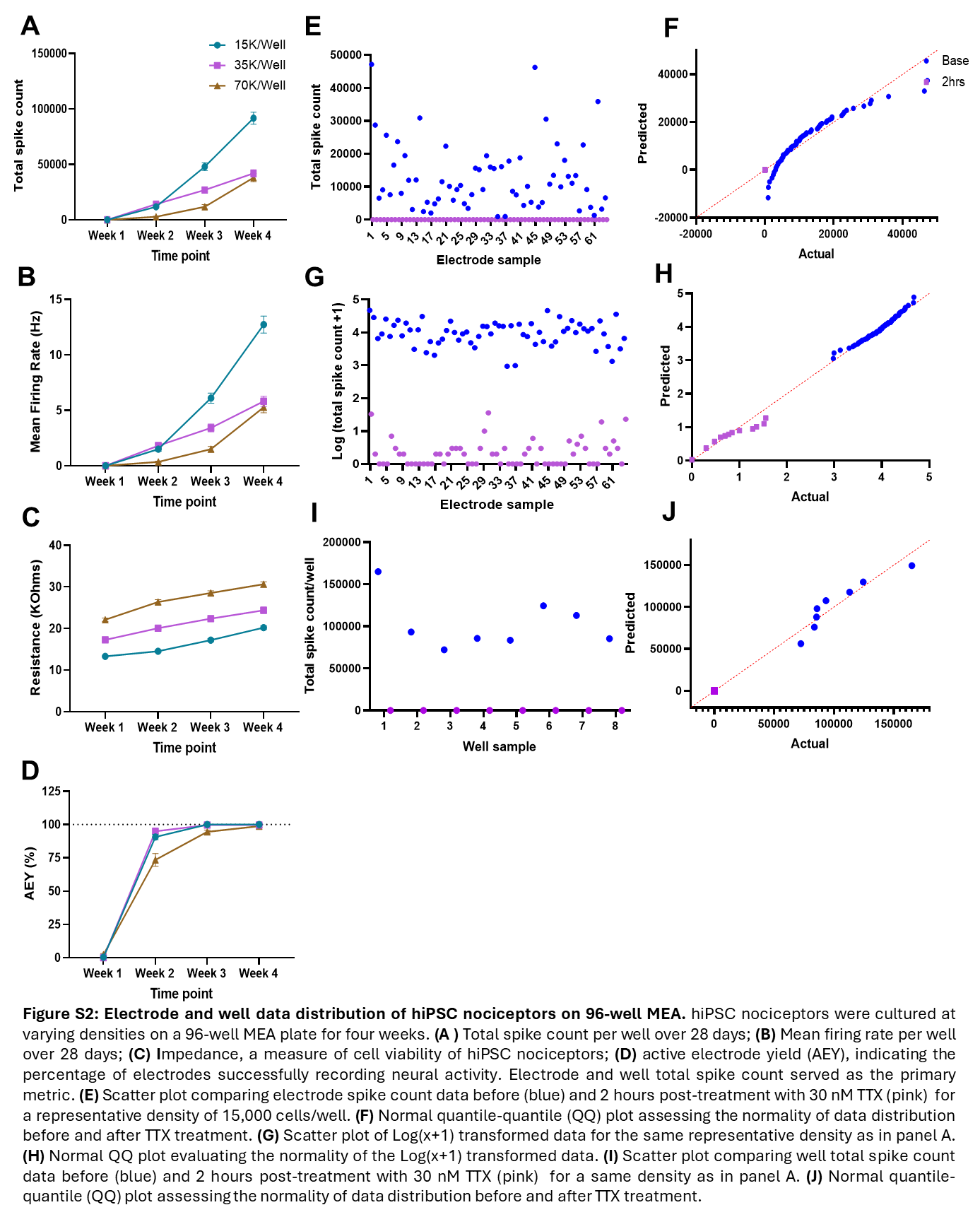

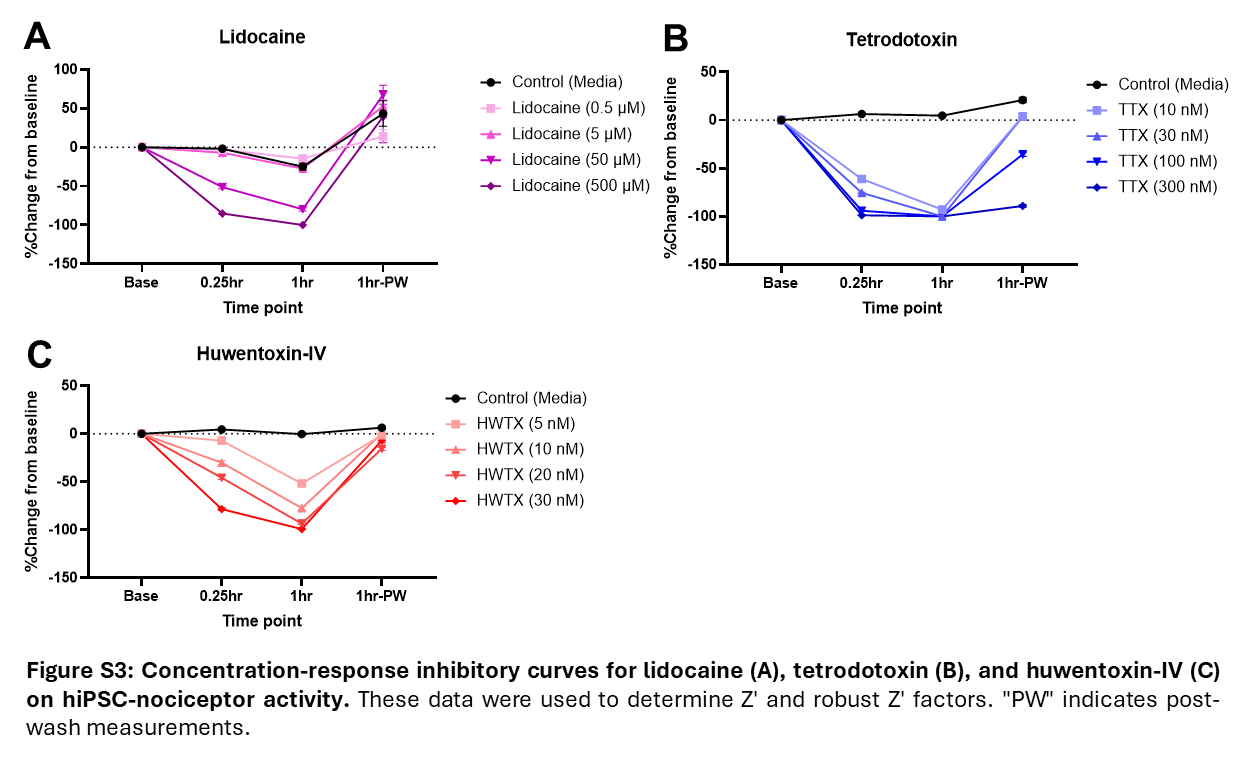

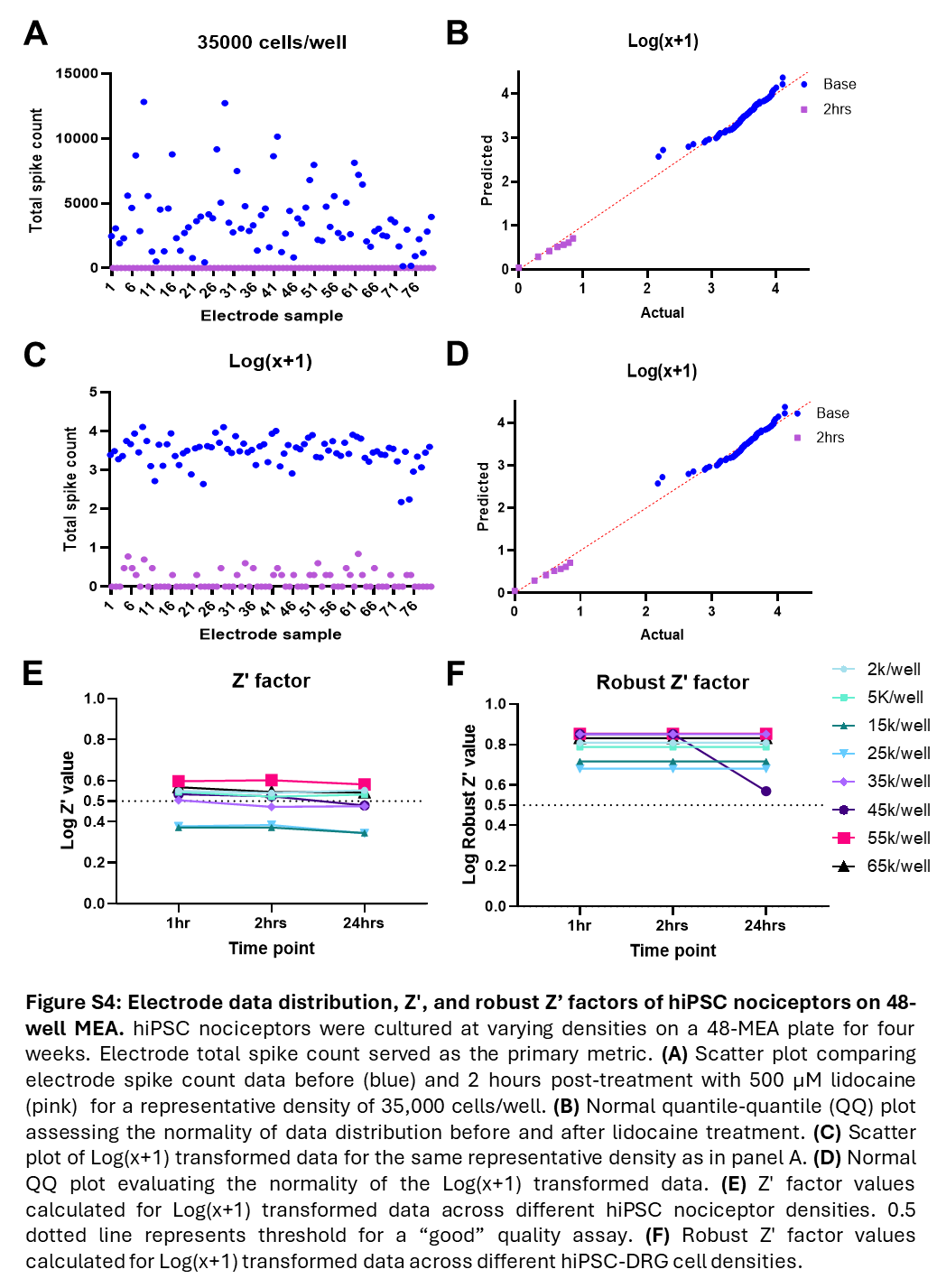

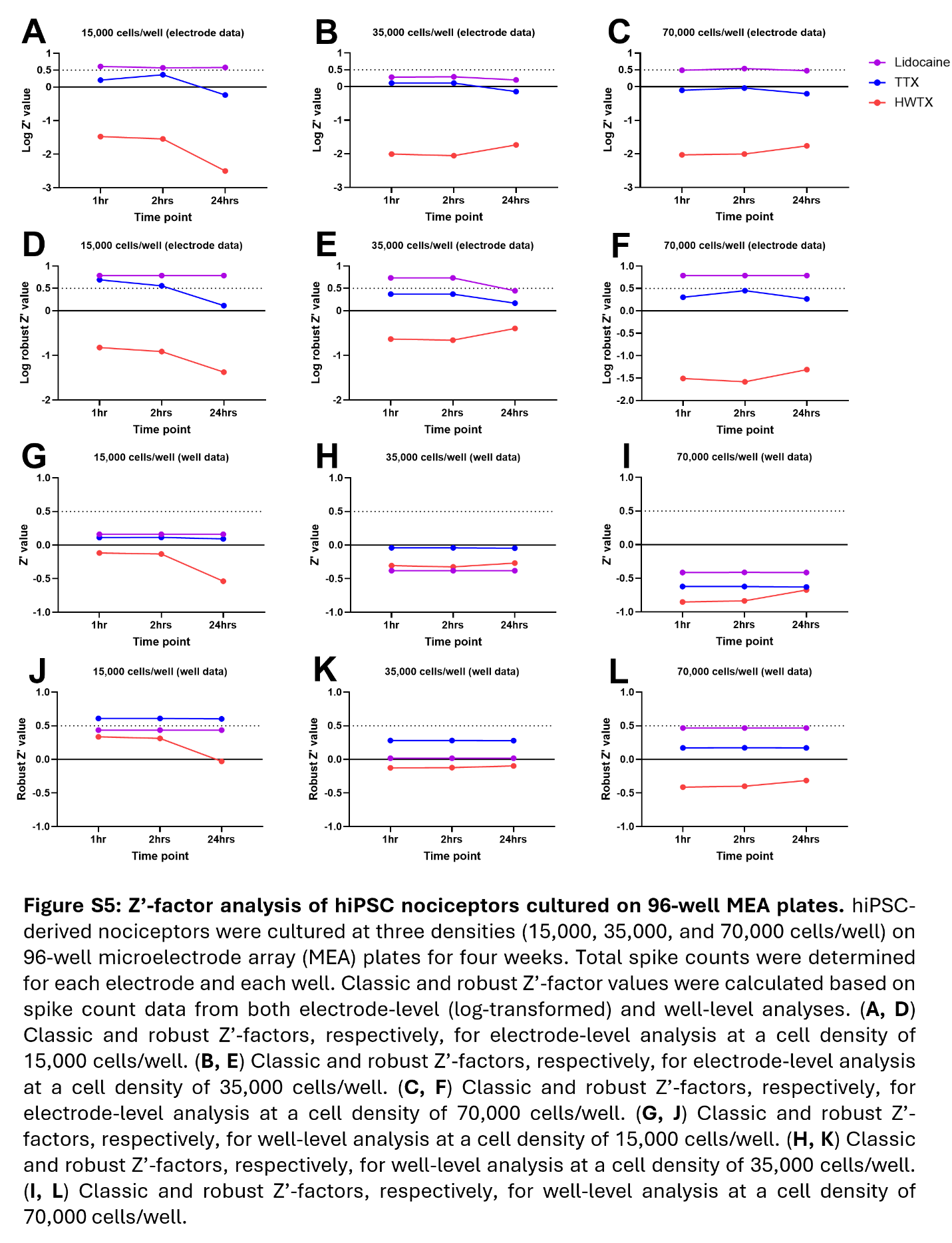

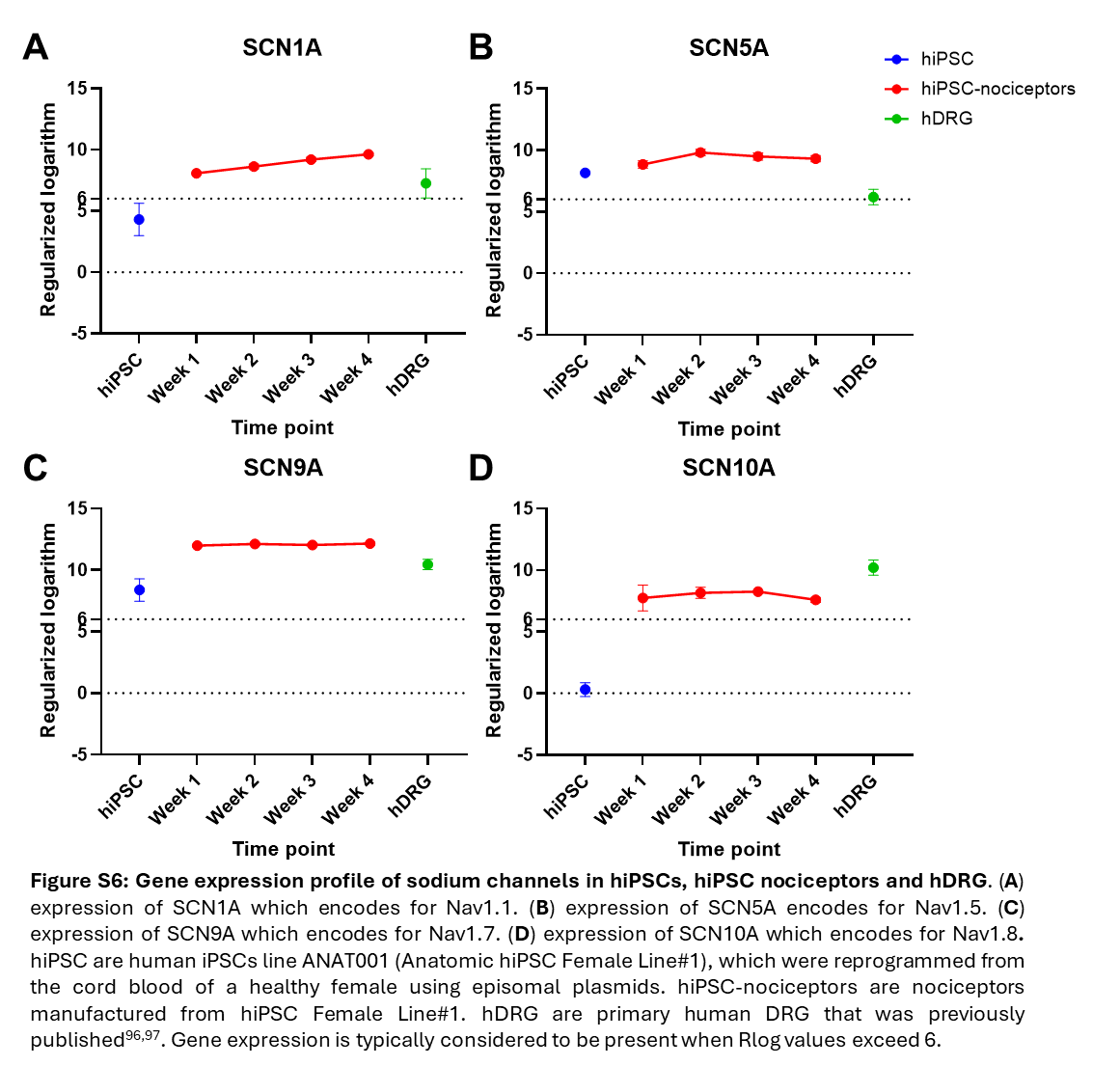

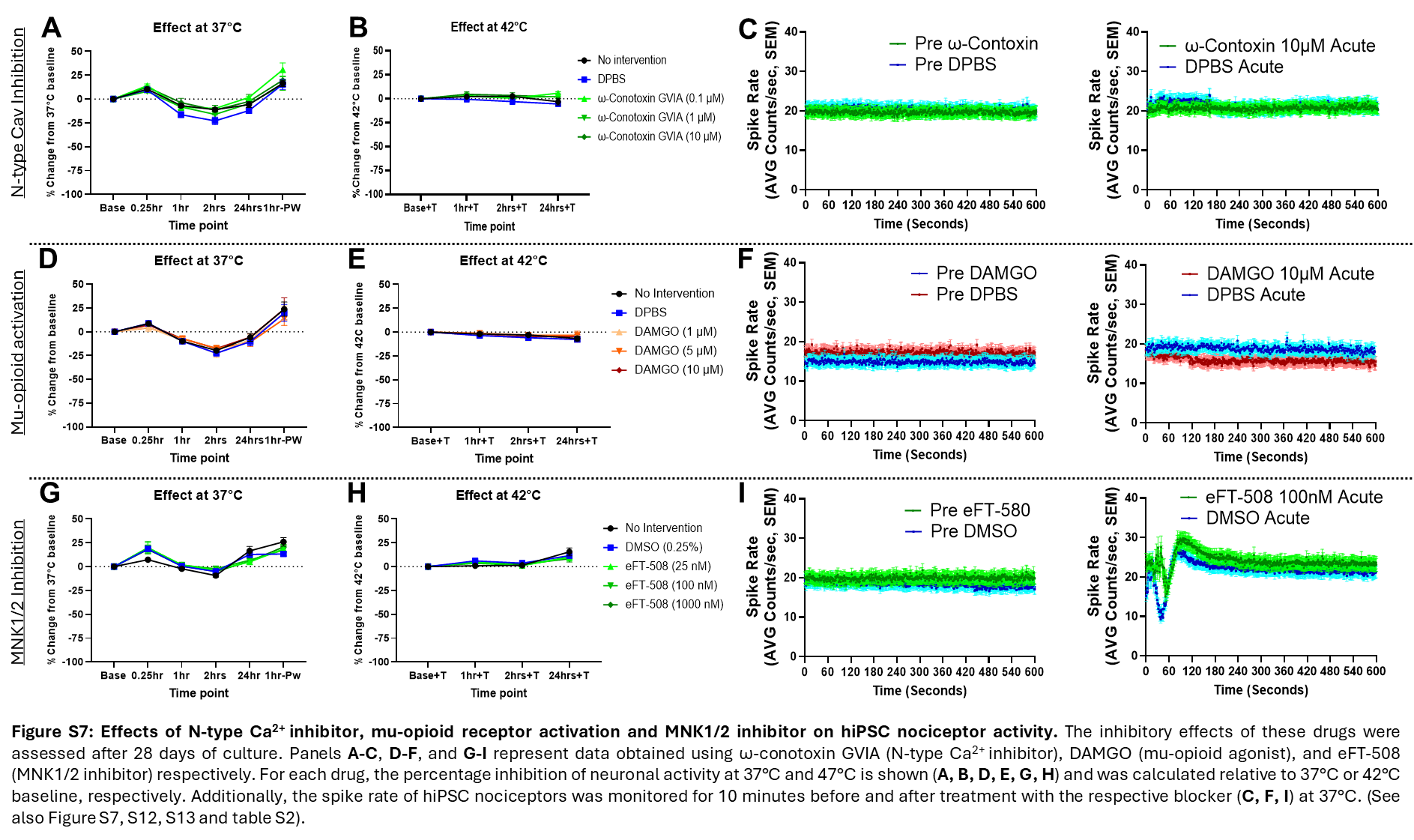

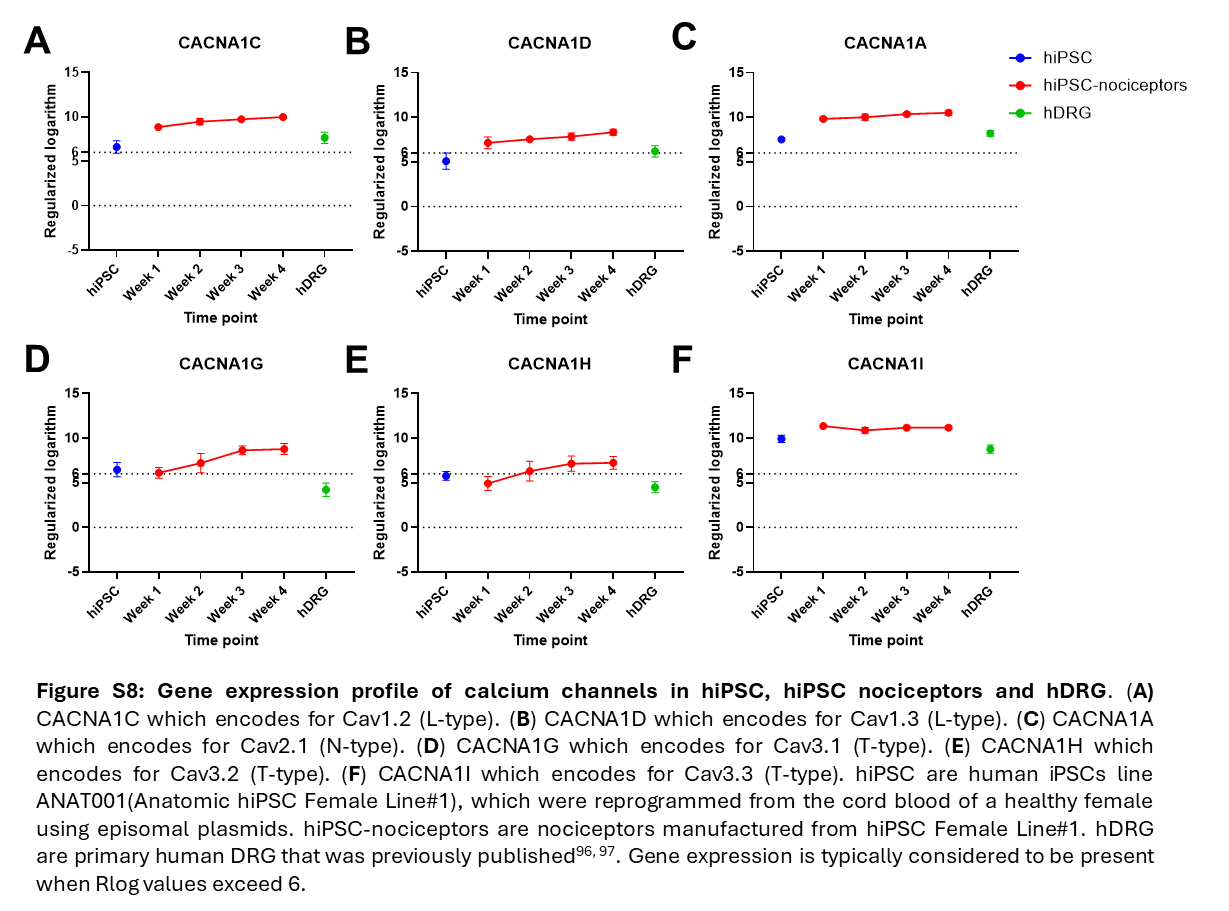

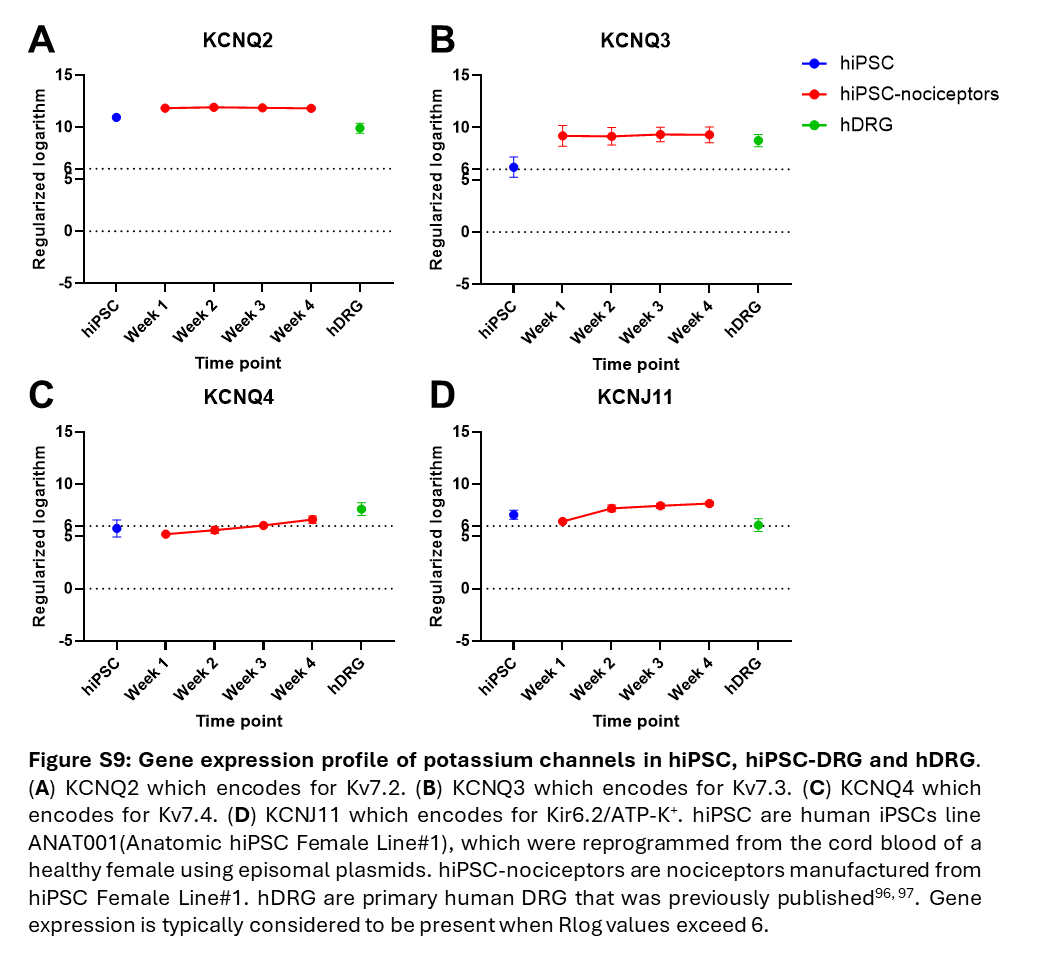

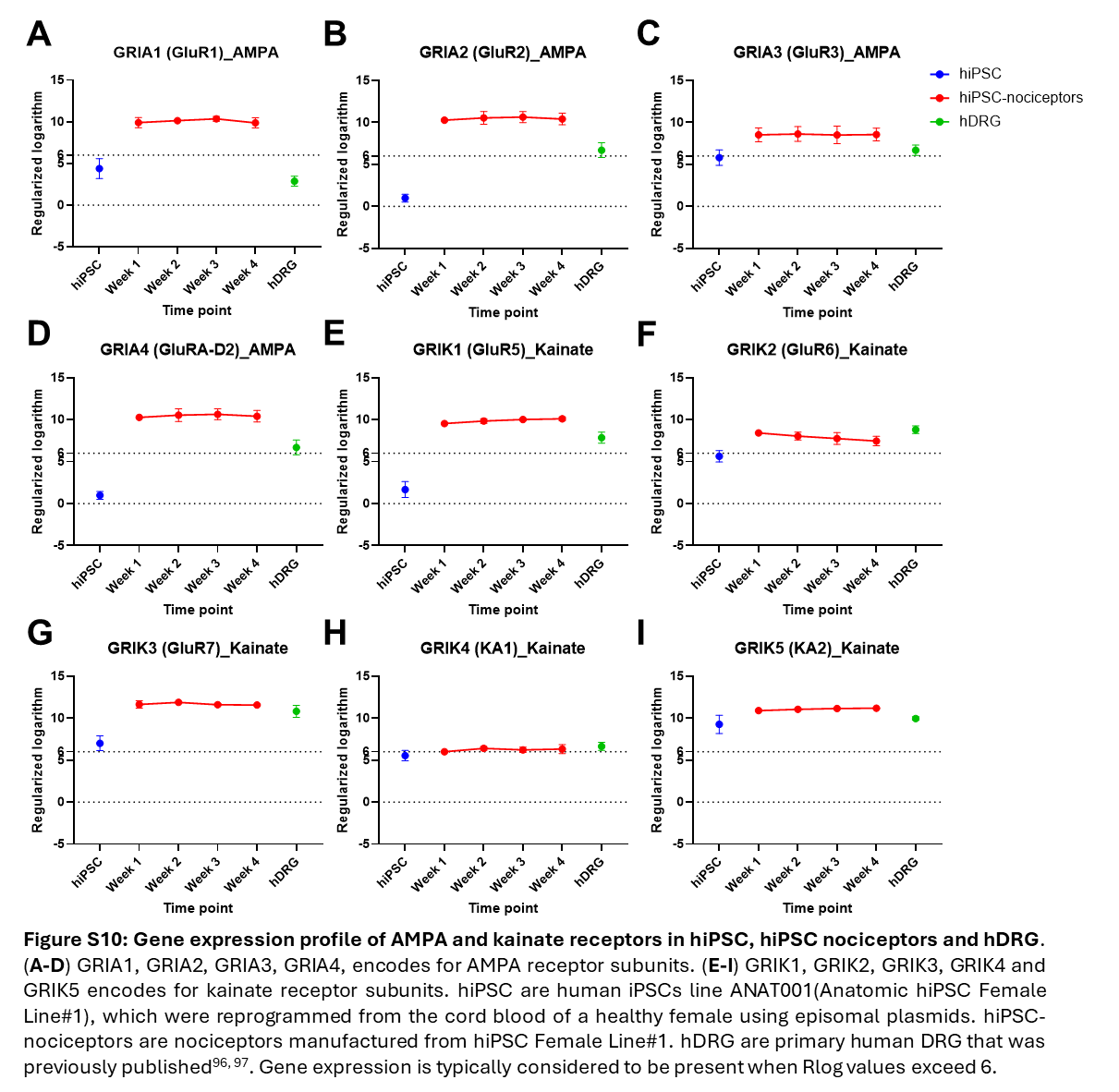

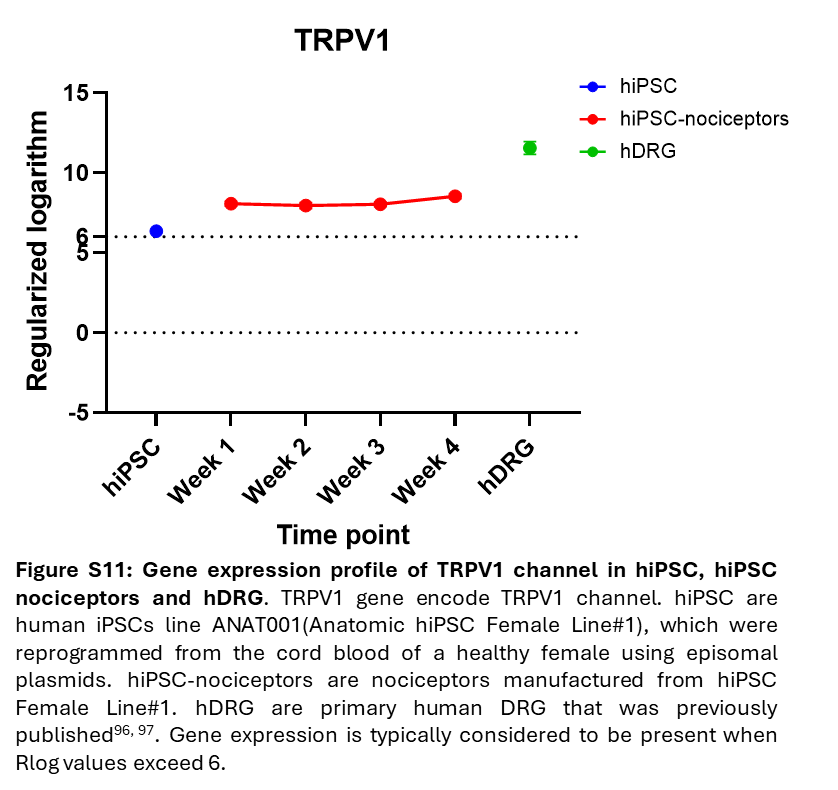

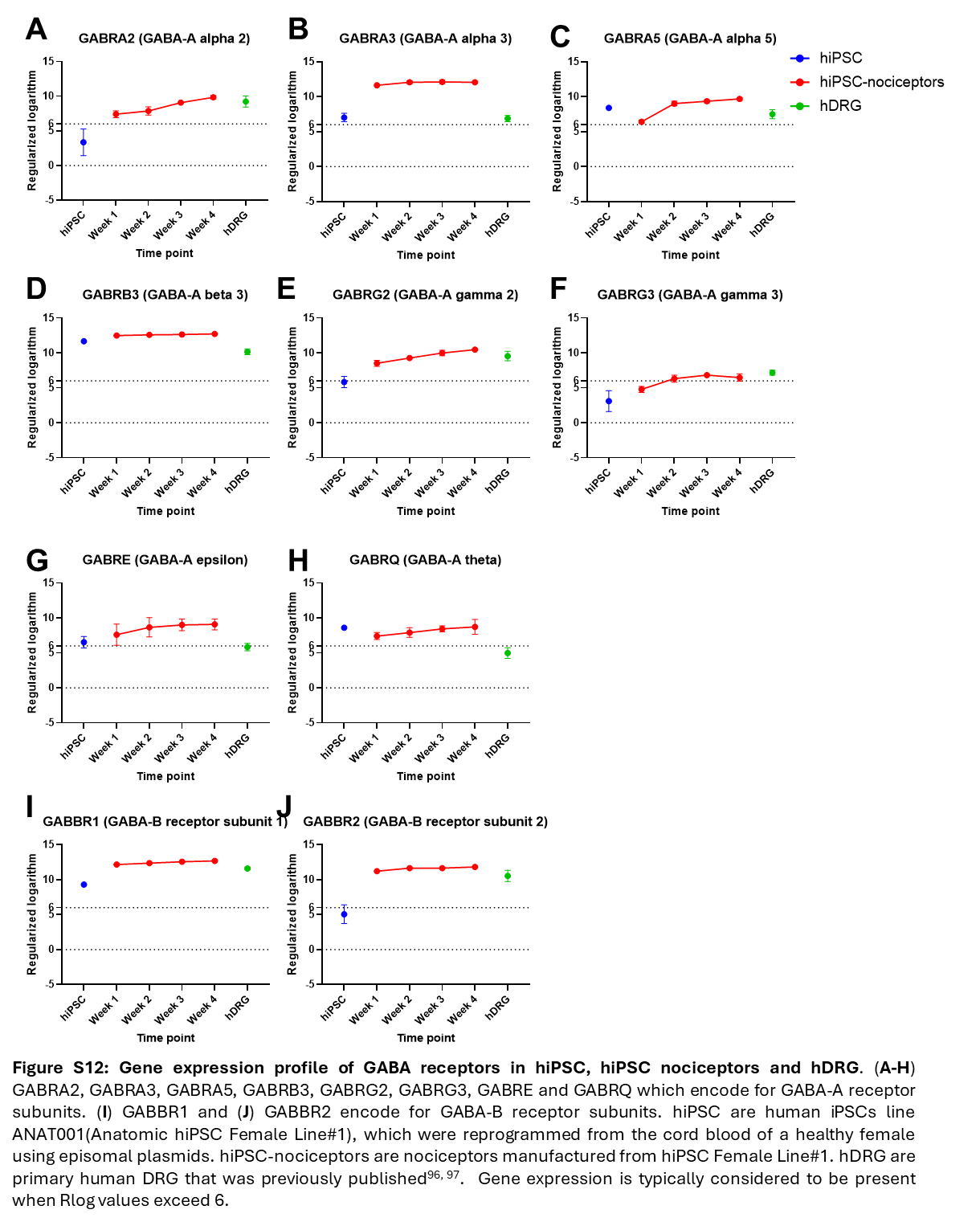

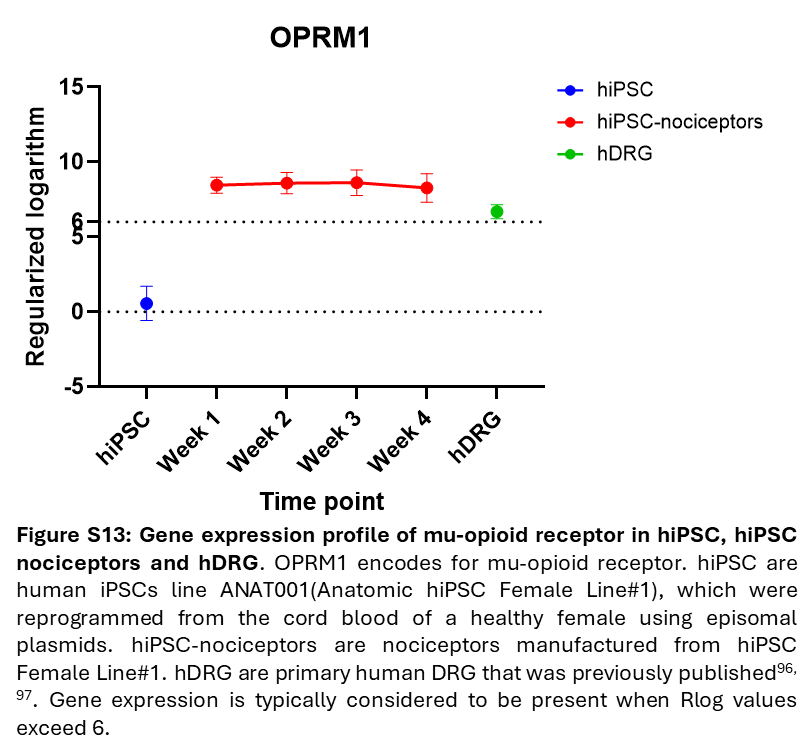

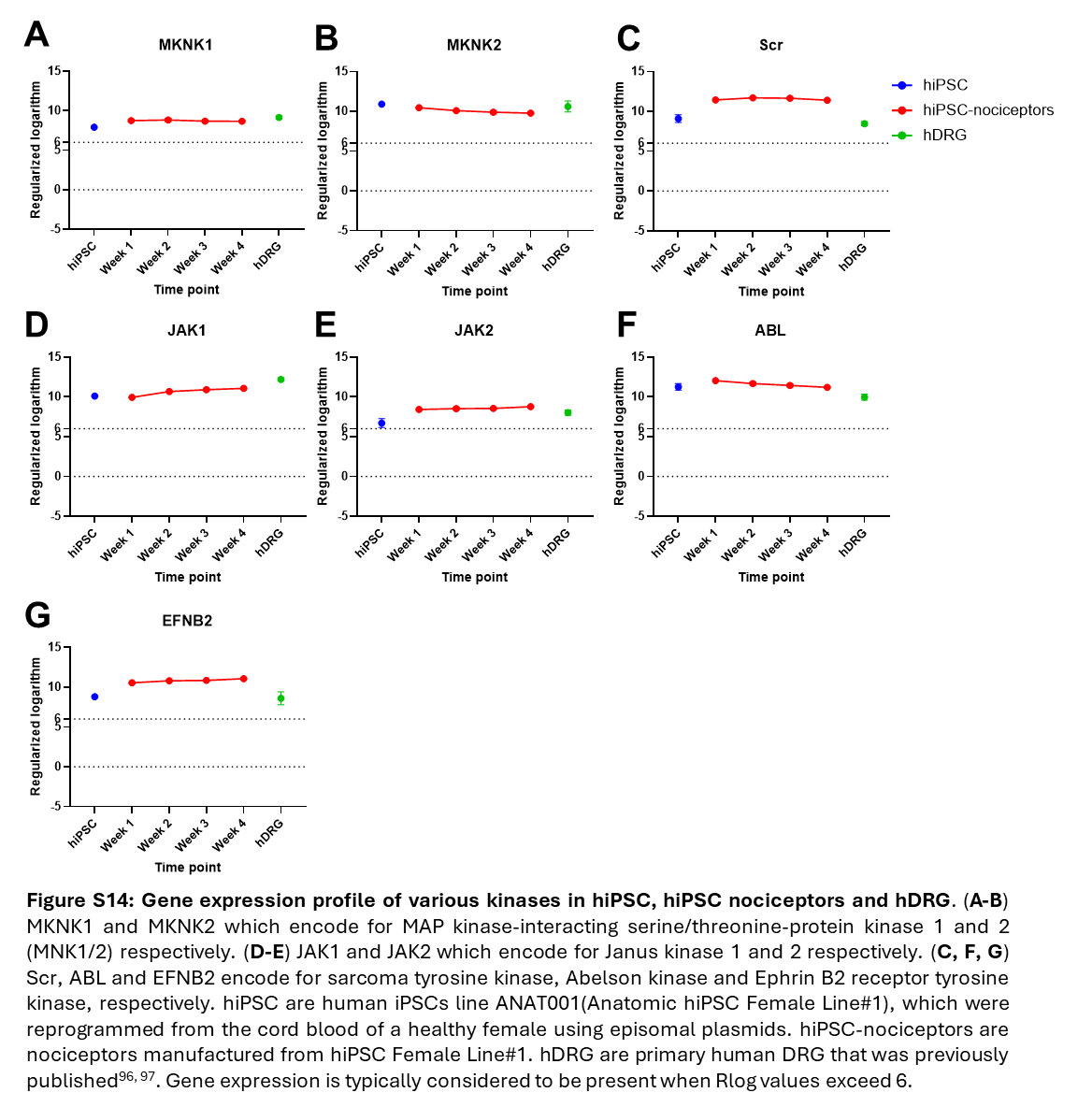
