## Supplemental Files Table S1-S3 for "Profiling Human iPSC-Derived Sensory Neurons for Analgesic Drug Screening Using a Multi-Electrode Array"

**Table S1:** Shapiro-Wilk test on data obtained from 96-well plate

|  | Shapiro-Wilk test on 96-well plate | | | | | |
| --- | --- | --- | --- | --- | --- | --- |
| Data | Electrode data | | Log transformed electrode data | | Well data | |
| Time point | Base | 2hrs | Base | 2hrs | Base | 2hrs |
| P value | <0.0001 | <0.0001 | 0.1955 | <0.0001 | 0.1169 | 0.0473 |
| Passed normality test (α = 0.05) | No | No | Yes | No | Yes | No |

**Table S2:** Adjusted p-values for pairwise comparisons of treatment groups to controls

| **Dunnett's multiple comparisons test** | **Adjusted P Value** | | | | | | | | | |
| --- | --- | --- | --- | --- | --- | --- | --- | --- | --- | --- |
|  | **Base-37°C** | **Base-42°C** | **0.25hr-37°C** | **1hr-**  **37°C** | **1hr-**  **42°C** | **2hr-**  **37°C** | **2hr-**  **42°C** | **24hr-**  **37°C** | **24hr-42°C** | **1hr-PW-37°C** |
| Control (DPBS) vs. No Intervention | 0 | 0 | 0.2951 | 0.8644 | 0.882 | 0.3355 | 0.5383 | >0.9999 | 0.6048 | 0.9886 |
| Control (DPBS) vs. TTX (5 nM) | 0 | 0 | 0.0065 | <0.0001 | <0.0001 | <0.0001 | <0.0001 | <0.0001 | <0.0001 | 0.996 |
| Control (DPBS) vs. TTX (15 nM) | 0 | 0 | <0.0001 | <0.0001 | <0.0001 | <0.0001 | <0.0001 | <0.0001 | <0.0001 | 0.0949 |
| Control (DPBS) vs. TTX (30 nM) | 0 | 0 | <0.0001 | <0.0001 | <0.0001 | <0.0001 | <0.0001 | <0.0001 | <0.0001 | 0.157 |
| Control (DPBS) vs. No Intervention | 0 | 0 | 0.2951 | 0.8644 | 0.882 | 0.3355 | 0.5383 | >0.9999 | 0.6048 | 0.9886 |
| Control (DPBS) vs. HWTX (5 nM) | 0 | 0 | >0.9999 | 0.9997 | 0.0956 | 0.9922 | 0.7818 | 0.9571 | 0.6013 | 0.0758 |
| Control (DPBS) vs. HWTX (10 nM) | 0 | 0 | 0.0378 | 0.0172 | 0.0008 | 0.0239 | 0.0047 | 0.0158 | 0.0012 | 0.1007 |
| Control (DPBS) vs. HWTX (20 nM) | 0 | 0 | <0.0001 | <0.0001 | <0.0001 | <0.0001 | <0.0001 | <0.0001 | <0.0001 | 0.0736 |
| DMSO (0.25%) vs. No Intervention | 0 | 0 | 0.4834 | 0.8617 | 0.0214 | 0.5035 | 0.3965 | >0.9999 | 0.9383 | 0.5919 |
| DMSO (0.25%) vs. GX201 (30 nM) | 0 | 0 | 0.6561 | 0.2567 | 0.0008 | 0.2059 | 0.0345 | 0.2086 | 0.1352 | 0.9996 |
| DMSO (0.25%) vs. GX201 (100 nM) | 0 | 0 | 0.0027 | 0.0232 | <0.0001 | 0.0216 | 0.0004 | 0.0074 | 0.0044 | >0.9999 |
| DMSO (0.25%) vs. GX201 (300 nM) | 0 | 0 | <0.0001 | 0.0002 | <0.0001 | 0.001 | 0.0002 | 0.0022 | 0.0004 | 0.0321 |
| DMSO (0.25%) vs. No Intervention | 0 | 0 | 0.1168 | 0.2955 | 0.421 | 0.4376 | 0.3965 | 0.0061 | 0.0448 | 0.3148 |
| DMSO (0.25%) vs. A887826 (0.1 μM) | 0 | 0 | 0.3838 | 0.2095 | 0.0029 | 0.3409 | 0.0068 | >0.9999 | 0.8911 | 0.9967 |
| DMSO (0.25%) vs. A887826 (0.5 μM) | 0 | 0 | 0.007 | 0.0014 | 0.0027 | 0.0011 | 0.0018 | 0.1245 | 0.0951 | 0.7909 |
| DMSO (0.25%) vs. A887826 (1 μM) | 0 | 0 | <0.0001 | <0.0001 | <0.0001 | <0.0001 | <0.0001 | <0.0001 | <0.0001 | 0.0044 |
| Control (DPBS) vs. No Intervention | 0 | 0 | 0.705 | 0.7279 | 0.9981 | 0.962 | >0.9999 | 0.9189 | 0.9967 | 0.7723 |
| Control (DPBS) vs. Mibefradil (0.5 μM) | 0 | 0 | 0.8656 | 0.7135 | 0.1715 | 0.3311 | 0.0862 | 0.767 | 0.5147 | 0.9833 |
| Control (DPBS) vs. Mibefradil (1 μM) | 0 | 0 | 0.7291 | 0.0408 | 0.0141 | 0.0003 | 0.0006 | 0.0003 | <0.0001 | 0.0001 |
| Control (DPBS) vs. Mibefradil (2 μM) | 0 | 0 | 0.0043 | <0.0001 | <0.0001 | <0.0001 | <0.0001 | <0.0001 | <0.0001 | <0.0001 |
| DPBS vs. No intervention | 0 | 0 | 0.9977 | 0.095 | 0.2243 | 0.0853 | 0.0573 | 0.4361 | 0.902 | 0.999 |
| DPBS vs. Conotoxin (0.1 μM) | 0 | 0 | 0.3956 | 0.2351 | 0.2434 | 0.0558 | 0.2288 | 0.011 | 0.0003 | 0.316 |
| DPBS vs. Conotoxin (1 μM) | 0 | 0 | 0.9011 | 0.3571 | 0.4252 | 0.6244 | 0.4264 | 0.5331 | 0.0633 | >0.9999 |
| DPBS vs. Conotoxin (10 μM) | 0 | 0 | 0.6192 | 0.1272 | 0.3141 | 0.2778 | 0.2406 | 0.6069 | 0.5349 | 0.9082 |
| DMSO (0.25%) vs. No intervention | 0 | 0 | 0.0065 | >0.9999 | 0.0384 | >0.9999 | 0.7825 | 0.9983 | 0.8535 | 0.4157 |
| DMSO (0.25%) vs. Nifedipine (10 µM) | 0 | 0 | 0.123 | 0.1029 | 0.9509 | 0.0582 | 0.3511 | 0.3712 | >0.9999 | >0.9999 |
| DMSO (0.25%) vs. Nifedipine (20 µM) | 0 | 0 | 0.0006 | 0.2051 | 0.7565 | 0.1012 | 0.8026 | 0.599 | 0.9935 | 0.9705 |
| DMSO (0.25%) vs. Nifedipine (30 µM) | 0 | 0 | <0.0001 | 0.0458 | 0.9592 | 0.0297 | 0.3116 | 0.3051 | 0.8619 | 0.8136 |
| Control (DPBS) vs. No Intervention | 0 | 0 | 0.705 | 0.7279 | 0.9981 | 0.962 | >0.9999 | 0.9189 | 0.9967 | 0.7723 |
| Control (DPBS) vs. Retigabine (2.5 µM) | 0 | 0 | 0.0021 | <0.0001 | <0.0001 | <0.0001 | <0.0001 | 0.0002 | 0.0001 | 0.9941 |
| Control (DPBS) vs. Retigabine (5 µM) | 0 | 0 | 0.0012 | <0.0001 | 0.0013 | <0.0001 | 0.0009 | <0.0001 | 0.0003 | 0.3953 |
| Control (DPBS) vs. Retigabine (10 µM) | 0 | 0 | <0.0001 | <0.0001 | <0.0001 | <0.0001 | <0.0001 | <0.0001 | <0.0001 | 0.9263 |
| DMSO (0.25%) vs. No intervention | 0 | 0 | 0.0065 | >0.9999 | 0.0384 | >0.9999 | 0.7825 | 0.9983 | 0.8535 | 0.4157 |
| DMSO (0.25%) vs. Diazoxide (25 µM) | 0 | 0 | <0.0001 | 0.3672 | 0.0376 | 0.2315 | 0.0844 | 0.911 | 0.4111 | 0.4049 |
| DMSO (0.25%) vs. Diazoxide (50 µM) | 0 | 0 | <0.0001 | 0.1334 | 0.0064 | 0.0725 | 0.0107 | 0.9953 | 0.6429 | 0.8763 |
| DMSO (0.25%) vs. Diazoxide (75 µM) | 0 | 0 | <0.0001 | 0.0017 | <0.0001 | 0.0019 | 0.0003 | 0.7377 | 0.248 | 0.9412 |
| DMSO (0.25%) vs. No Intervention | 0 | 0 | 0.1168 | 0.2955 | 0.421 | 0.4376 | 0.3965 | 0.0061 | 0.0448 | 0.3148 |
| DMSO (0.25%) vs. DNQX (50 μM) | 0 | 0 | <0.0001 | 0.0165 | 0.2747 | 0.0127 | 0.2538 | 0.6317 | 0.994 | 0.9667 |
| DMSO (0.25%) vs. DNQX (75 μM) | 0 | 0 | <0.0001 | 0.0008 | 0.0202 | <0.0001 | 0.0157 | 0.1204 | 0.9946 | 0.332 |
| DMSO (0.25%) vs. DNQX (100 μM) | 0 | 0 | <0.0001 | <0.0001 | <0.0001 | <0.0001 | <0.0001 | 0.0004 | 0.1933 | 0.4093 |
| DMSO (0.25%) vs. No Intervention | 0 | 0 | 0.0252 | 0.9704 | 0.0569 | 0.9674 | 0.7332 | 0.1963 | 0.1495 | 0.7464 |
| DMSO (0.25%) vs. Capsazepine (10 μM) | 0 | 0 | <0.0001 | 0.0185 | 0.006 | <0.0001 | <0.0001 | <0.0001 | <0.0001 | 0.2069 |
| DMSO (0.25%) vs. Capsazepine (20 μM) | 0 | 0 | <0.0001 | 0.001 | 0.0005 | <0.0001 | 0.0005 | <0.0001 | 0.0006 | 0.2342 |
| DMSO (0.25%) vs. Capsazepine (30 μM) | 0 | 0 | <0.0001 | <0.0001 | <0.0001 | <0.0001 | <0.0001 | <0.0001 | <0.0001 | 0.0014 |
| DMSO (0.25%) vs. No Intervention | 0 | 0 | 0.0396 | 0.6682 | 0.0762 | 0.3837 | 0.3705 | 0.901 | 0.8152 | 0.1159 |
| DMSO (0.25%) vs. eFT-508 (25 nM) | 0 | 0 | 0.9985 | 0.9416 | 0.9788 | 0.9438 | 0.4506 | 0.247 | 0.9865 | 0.6314 |
| DMSO (0.25%) vs. eFT-508 (100 nM) | 0 | 0 | 0.9988 | 0.9741 | 0.7237 | 0.6826 | 0.4857 | 0.4096 | 0.9863 | 0.9101 |
| DMSO (0.25%) vs. eFT-508 (1000 nM) | 0 | 0 | >0.9999 | 0.9996 | 0.9185 | >0.9999 | 0.7056 | 0.3604 | 0.8668 | 0.5603 |
| DMSO (0.25%) vs. No Intervention | 0 | 0 | 0.0252 | 0.9704 | 0.0569 | 0.9674 | 0.7332 | 0.1963 | 0.1495 | 0.7464 |
| DMSO (0.25%) vs. Dasatinib (5 μM) | 0 | 0 | <0.0001 | 0.0135 | 0.0025 | 0.0032 | 0.0074 | 0.9965 | 0.9759 | 0.2643 |
| DMSO (0.25%) vs. Dasatinib (10 μM) | 0 | 0 | <0.0001 | 0.0077 | 0.0037 | 0.0082 | 0.0376 | 0.9681 | 0.9058 | 0.9925 |
| DMSO (0.25%) vs. Dasatinib (20 μM) | 0 | 0 | <0.0001 | <0.0001 | <0.0001 | <0.0001 | <0.0001 | <0.0001 | 0.0002 | 0.0982 |
| DMSO (0.25%) vs. No Intervention | 0 | 0 | 0.213 | 0.9991 | 0.0914 | 0.6331 | 0.9236 | 0.0971 | 0.0693 | 0.602 |
| DMSO (0.25%) vs. Baricitinib (5 μM) | 0 | 0 | <0.0001 | 0.4517 | 0.0176 | 0.4254 | 0.1574 | 0.0273 | 0.0962 | 0.9321 |
| DMSO (0.25%) vs. Baricitinib (10 μM) | 0 | 0 | <0.0001 | 0.1208 | 0.001 | 0.0132 | 0.0035 | 0.0007 | 0.004 | 0.2191 |
| DMSO (0.25%) vs. Baricitinib (20 μM) | 0 | 0 | 0.0003 | 0.0397 | 0.3018 | 0.076 | 0.2075 | 0.0053 | 0.0487 | 0.1664 |
| DPBS vs. No Intervention | 0 | 0 | 0.9999 | >0.9999 | 0.7435 | 0.6026 | 0.589 | 0.8746 | 0.9958 | 0.9901 |
| DPBS vs. DAMGO (1 μM) | 0 | 0 | 0.7391 | 0.9944 | 0.9998 | >0.9999 | >0.9999 | 0.9908 | >0.9999 | >0.9999 |
| DPBS vs. DAMGO (5 μM) | 0 | 0 | 0.4374 | 0.6195 | 0.4989 | 0.497 | 0.7107 | >0.9999 | 0.7874 | 0.9452 |
| DPBS vs. DAMGO (10 μM) | 0 | 0 | 0.8654 | 0.4979 | 0.9995 | 0.2893 | 0.9789 | 0.8125 | 0.7254 | 0.9947 |
| DPBS vs. No intervention | 0 | 0 | 0.9977 | 0.095 | 0.2243 | 0.0853 | 0.0573 | 0.4361 | 0.902 | 0.999 |
| DPBS vs. GABA (10 μM) | 0 | 0 | 0.0112 | 0.0908 | 0.8445 | 0.9998 | 0.9429 | 0.0457 | 0.0481 | 0.9995 |
| DPBS vs. GABA (50 μM) | 0 | 0 | <0.0001 | <0.0001 | 0.0001 | <0.0001 | <0.0001 | 0.0082 | 0.9992 | 0.7499 |
| DPBS vs. GABA (100 μM) | 0 | 0 | <0.0001 | <0.0001 | 0.0016 | <0.0001 | 0.0011 | 0.0004 | 0.363 | 0.8974 |
| DMSO (0.25%) vs. No Intervention | 0 | 0 | 0.213 | 0.9991 | 0.0914 | 0.6331 | 0.9236 | 0.0971 | 0.0693 | 0.602 |
| DMSO (0.25%) vs. Duloxetine (0.5 μM) | 0 | 0 | 0.0233 | 0.2361 | 0.0005 | 0.0533 | 0.033 | 0.1041 | 0.0146 | 0.5819 |
| DMSO (0.25%) vs. Duloxetine (1 μM) | 0 | 0 | <0.0001 | 0.0015 | <0.0001 | 0.0003 | <0.0001 | 0.0055 | 0.0007 | 0.0277 |
| DMSO (0.25%) vs. Duloxetine (5 μM) | 0 | 0 | <0.0001 | <0.0001 | <0.0001 | <0.0001 | <0.0001 | <0.0001 | <0.0001 | 0.0001 |
| DMSO (0.25%) vs. No Intervention | 0 | 0 | 0.4834 | 0.8617 | 0.0214 | 0.5035 | 0.3965 | >0.9999 | 0.9383 | 0.5919 |
| DMSO (0.25%) vs. Amitriptyline (0.5 μM) | 0 | 0 | <0.0001 | 0.001 | <0.0001 | 0.0017 | 0.0002 | 0.0027 | 0.0046 | 0.847 |
| DMSO (0.25%) vs. Amitriptyline (0.75 μM) | 0 | 0 | <0.0001 | 0.0002 | 0.0004 | 0.0003 | <0.0001 | 0.0002 | 0.0004 | 0.1124 |
| DMSO (0.25%) vs. Amitriptyline (1 μM) | 0 | 0 | <0.0001 | <0.0001 | <0.0001 | 0.0002 | <0.0001 | <0.0001 | <0.0001 | 0.1248 |

This table presents the adjusted P-values from Dunnett's multiple comparisons test, comparing various experimental treatments to control groups (DPBS or DMSO) across different time points (0, 0.25, 1, 2, and 24 hours) and temperatures (37°C and 42°C). Treatments included TTX, HWTX, GX201, A887826, Mibefradil, Retigabine, Diazoxide, DNQX, Capsazepine, eFT-508, Dasatinib, Baricitinib, DAMGO, GABA, Duloxetine, and Amitriptyline. P-values less than 0.05 indicate statistically significant differences between the treatment group and the respective control.

**Table S3: Reagents and resources**

| **Item** | **Catalogue number** | **Company** | **Description** |
| --- | --- | --- | --- |
| Maestro Pro |  | Axion | Recording device |
| 48-well MEA plate | M768-tMEA-48W | Axion | MutiWell Electrode Array |
| 96-well MEA plate | M768-tMEA-96W | Axion | MutiWell Electrode Array |
| Poly-L-Ornithine | A-004-C | EMD Millipore Sigma | Coating |
| iMatrix-511 | SKU:N-892022 | Iwai-chem | Coating |
| dPBS | D8537-100ML | Sigma-Aldrich |  |
| DMEM/F12 | 11320-033 | Gibco™ | Culture Media |
| Anatomic Senso-MM | 1030 | Anatomic | Culture Media |
| Lidocaine | J63035.06 | Thermo Scientific | Nav inhibitor |
| TTX | 1078/1 | Tocris | Nav1.7 inhibitor |
| Huwentoxin IV | STH-100 | Alomone Labs | Nav1.7 inhibitor |
| DMSO | 67-68-5 | Fisher Scientific | Solvent |
| GX201 | **7029/10** | Tocris | Nav1.7 inhibitor |
| A-887826 | 4249 | Tocris | Nav1.8 inhibitor |
| Retigabine | A3758 | APExBIO | KCNQ1-5/Kv7 opener |
| diazoxide | A601456 | AmBeed | K_ATP_ activator |
| Nifedipine | A516303 | AmBeed | L-type calcium channel blocker |
| ω-conotoxin GVIA | C-300 | Alomone labs | N-type calcium channel blocker |
| Mibefradil | M5441-5MG | Sigma-Aldrich | T-type calcium channel blocker |
| GABA | A2129 | Sigma-Aldrich | GABA_A/B_ agonist |
| eFT-508 | S8275 | Selleckchem | MNK inhibitor |
| Dasatinib | A355193 | AmBeed | *pan*-tyrosine kinase inhibitor |
| Baricitinib | A214136 | AmBeed | janus kinase, JAK1/2 inhibitor |
| DNQX | D0540 | Sigma-Aldrich | AMPA, kainite antagonist |
| D-(-)APV | A8054 | Sigma-Aldrich | NMDA antagonist |
| Capsazepine | C-120 | Alomone labs | TRPV1 antagonist |
| Duloxetine | CS-1993 | Chem Scene | SNRI |
| Amitriptyline | 37560 | Chem-Impex | SNRI |
| DAMGO | HY-P0210 | MedChemExpress | mu-opioid agonist |
